# Genetic analysis reveals a robust and hierarchical recruitment of the LolA chaperone to the LolCDE lipoprotein transporter

**DOI:** 10.1101/2023.11.08.566237

**Authors:** Kelly M. Lehman, Kerrie L. May, Julianna Marotta, Marcin Grabowicz

## Abstract

The outer membrane (OM) is an essential organelle of Gram-negative bacteria. Lipoproteins are key to building the OM, performing essential functions in several OM assembly machines. Lipoproteins mature in the inner membrane (IM) and are then trafficked to the OM. In *Escherichia coli*, the LolCDE transporter is needed to extract lipoproteins from the IM to begin trafficking. Lipoproteins are then transferred from LolCDE to the periplasmic chaperone LolA which ferries them to the OM for insertion by LolB. LolA recruitment by LolC is an essential trafficking step. Structural and biochemical studies suggested that two regions (termed Hook and Pad) within a periplasmic loop of LolC worked in tandem to recruit LolA, leading to a bipartite model for recruitment. Here, we genetically examine the LolC periplasmic loop *in vivo* using *E. coli*. Our findings challenge the bipartite interaction model. We show that while the Hook is essential for lipoprotein trafficking *in vivo*, lipoproteins are still efficiently trafficked when the Pad residues are inactivated. We show with AlphaFold2 multimer modeling that Hook:LolA interactions are likely universal among diverse Gram-negative bacteria. Conversely, Pad:LolA interactions vary across phyla. Our *in vivo* data redefine LolC:LolA recruitment into a hierarchical interaction model. We propose that the Hook is the major player in LolA recruitment, while the Pad plays an ancillary role that is important for efficiency but is ultimately dispensable. Our findings expand the understanding of a fundamental step in essential lipoprotein trafficking and have implications for efforts to develop new antibacterials that target LolCDE.

**IMPORTANCE:** Resistance to current antibiotics is increasingly common. New antibiotics that target essential processes are needed to expand clinical options. For Gram-negative bacteria, their cell surface— the outer membrane (OM)—is an essential organelle and antibiotic barrier that is an attractive target for new antibacterials. Lipoproteins are key to building the OM. The LolCDE transporter is needed to supply the OM with lipoproteins and has been a focus of recent antibiotic discovery. *In vitro* evidence recently proposed a two-part interaction of LolC with LolA lipoprotein chaperone (which traffics lipoproteins to the OM) via “Hook” and “Pad” regions. We show that this model does not reflect lipoprotein trafficking *in vivo*. Only the Hook is essential for lipoprotein trafficking and is remarkably robust to mutational changes. The Pad is non-essential for lipoprotein trafficking but plays an ancillary role, contributing to trafficking efficiency. These insights inform ongoing efforts to drug LolCDE.

## INTRODUCTION

For Gram-negative bacteria (like *Escherichia coli*), the outer membrane (OM) is an essential organelle (1, 2). The OM also serves as a barrier opposing the entry of toxic molecules, including many antibiotics (3, 4). The barrier property of the OM limits clinical options for treating Gram-negative infections, especially as resistance to current antibiotics continues to spread. Indeed, *E. coli* is globally responsible for the most deaths directly attributable to antibiotic resistance, and four of the top six deadly pathogens are Gram-negative species (5). Efforts to develop new antibiotics have included a focus on disrupting OM assembly (6). Several conserved and essential molecular machines work to assemble the OM (8–10). Most of these machines require one or more OM lipoproteins to function (10). Preventing lipoproteins from reaching these OM machines represents a key vulnerability that can be therapeutically exploited. Lipoproteins are a family of proteins that are triacylated in the inner membrane (IM) following their secretion from the cytosol (11, 12). Most lipoproteins in *E. coli* are targeted to the OM and must therefore traverse the aqueous periplasm. The localization <u>o</u>f lipoproteins (Lol) pathway is the canonical pathway responsible for trafficking lipoproteins from the IM, across the periplasm, and delivering them to the OM (**Fig. 1A**) (10, 13). The first step in the *E. coli* Lol pathway relies on an ATP-binding cassette (ABC) transporter consisting of a transmembrane protein heterodimer of LolC and LolE that is coupled to a homodimer LolD ATPase (14–16). The LolCDE transporter extracts lipoproteins from the IM and transfers them to the periplasmic chaperone LolA (17). Cargo-laden LolA then delivers lipoproteins to LolB at the OM for insertion into the bilayer (18). While LolAB are the canonical trafficking route, they are dispensable in *E. coli* under defined genetic conditions, implying the existence of an alternate trafficking route (19). LolCDE is essential whether lipoprotein trafficking proceeds through either LolAB or alternate routes, demonstrating that LolCDE is the origin of all lipoproteins destined for the OM (19).

**Figure 1.**
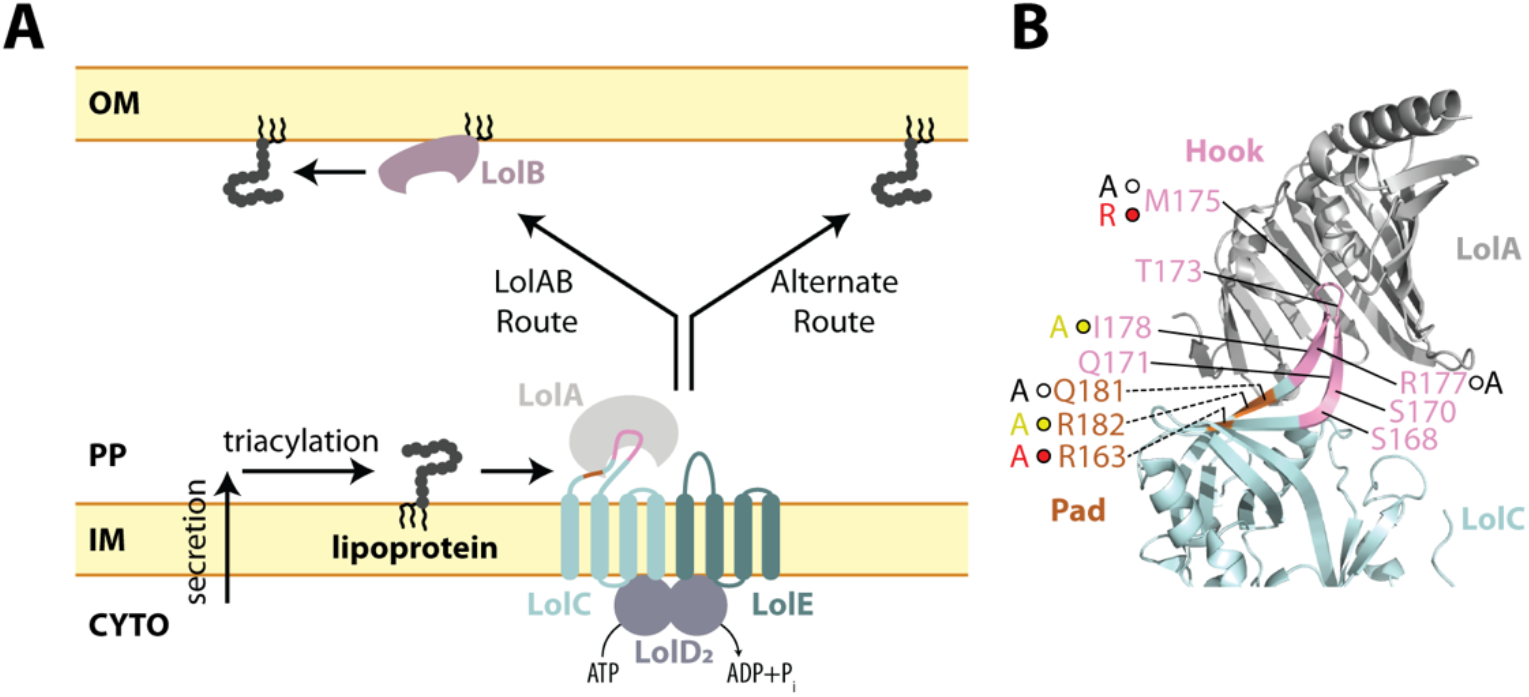
The Lol lipoprotein trafficking pathway of *E. coli* and LolC:LolA interaction. **(A)** Secreted and triacylated lipoproteins can enter the OM trafficking pathway. The LolCDE transporter uses cycles of ATP hydrolysis to extract lipoproteins from the IM, transfer them to LolA, and reset for the next lipoprotein. LolA and LolB form one trafficking route to the OM and in their absence an alternate route delivers lipoproteins to the OM. LolCDE is essential for all trafficking routes. **(B)** Summary of LolC mutations constructed in this study on the structure of LolA with the periplasmic domain of LolC (PDB 6F3Z). Direction of labeling lines indicates orientation of side chains. The Hook region (aa 167-179) and Pad residues (aa 163, 181, and 182) in LolC are colored pink and orange, respectively. Mutations that were previously tested biochemically are shown with spheres and the changed residues. Spheres indicate severity of defect that mutations cause to the biochemical interaction of LolC:LolA reported by Kaplan *et al*.: red spheres indicate non-binding variants, yellow spheres indicate severely defective binders, and white spheres indicate modestly defective binders (25). The amino acid substitution for each Kaplan *et al*. mutation is provided next to each sphere (25). Residues lacking spheres were additionally examined by mutagenesis in the current study.

The pivotal role of LolCDE in OM lipoprotein trafficking has made it an attractive target for novel antibiotics. Indeed, a CRISPRi screen of all essential *Vibrio cholerae* genes suggested that this organism is most acutely sensitive to depletion of LolC levels (20). Recent industry efforts have independently identified several compounds that can inhibit LolCDE activity (21–23). Nonetheless, a detailed understanding of how LolCDE functions is needed to refine therapeutic efforts targeting this essential transporter.

LolCDE is an unusual ABC transporter since it does not import or export molecules across the IM. Instead, LolCDE extracts lipoproteins from the IM bilayer for subsequent trafficking to the OM. This activity is similar to that of the LptBFG ABC transporter which extracts lipopolysaccharide (LPS) molecules from the IM for transport to the OM via the LptA transenvelope bridge (8, 24). However, unlike LptBFG, which is thought to form a stable complex with the LptA bridge via an interaction with LptC, LolCDE must dynamically cycle between recruiting the periplasmic chaperone LolA, loading it with lipoprotein cargo, and then releasing it.

Insights into how LolA is recruited to the LolCDE complex have only recently begun to emerge from studies of the *E. coli* system (25). By co-purifying the large periplasmic loop of LolC and mixing it with purified LolA, Kaplan *et al.* were able to crystalize the interaction of this loop with LolA (25). Two regions—Hook and Pad—within the loop appeared to be involved in the LolC:LolA interaction (25). While both LolC and LolE have a large periplasmic domain, only the LolC domain could bind LolA *in vitro*, implying that specific residues present in LolC (but absent in LolE) mediate LolA recruitment (25). Indeed, many LolC substitutions appeared to strongly disrupt or entirely abolish LolC:LolA interactions *in vitro* as judged by gel filtration assays (summarized in **Fig. 1B**) (25). A cryo-EM structural study independently supported that these interactions likely occur in the context of both full-length LolC and the larger LolCDE complex (26). Yet, the *in vivo* significance of the LolC Hook or Pad for lipoprotein trafficking has not been investigated. Moreover, whether the LolC:LolA interaction described in the *E. coli* system is widely conserved among Gram-negative bacteria is not known. Key differences in lipoprotein trafficking exist outside of *E. coli* and its closely related bacterial cousins: while the *E. coli* transporter consists of a LolC-LolE heterodimeric transmembrane protein complex, many Proteobacteria (the major Gram-negative phylum) produce a homodimeric transmembrane protein complex consisting of a LolF hybrid protein (27).

In this study, we use AlphaFold2 Multimer modeling to predict how LolA recruitment by LoC/LolF among diverse bacterial species is likely to occur. We find that LolA proteins are predicted to be recruited by the Hook in all our models. Using genetic analysis of *lolC* in *E. coli*, we show that the Hook region is fundamentally essential, in agreement with biochemical evidence that this region is needed for LolA recruitment. Surprisingly, however, we also find that the Hook is remarkably permissive for mutations. LolC substitutions that abolished or severely weakened the LolC:LolA interaction in biochemical assays are robustly tolerated *in vivo*. Importantly, we demonstrate that the Pad is not essential for lipoprotein trafficking *in vivo* but does contribute to efficient trafficking. Our findings place structural insights into biological context and redefine the mechanism of how LolC recruits LolA. Rather than a bipartite Hook and Pad model, we reveal a hierarchical recruitment mechanism in which the Hook is the primary, essential, and likely conserved recruiter of LolA. Meanwhile, we show that the Pad contributes to increasing the efficiency of trafficking, rather than contributing to a fundamentally essential step in trafficking.

## RESULTS

### Conservation of Hook and Pad interactions with LolA among diverse Proteobacteria

The current understanding of LolA recruitment to the lipoprotein trafficking transporter (LolCDE or LolDF) is based on structural and biochemical analysis of the *E. coli* system (25, 26). Whether this paradigm is applicable to evolutionarily distant relatives has not been explored. To gain insights into how LolA might be recruited among diverse species, we selected paired LolA and LolC/LolF sequences from representative species across the Proteobacterial phylum and used AlphaFold2 Multimer prediction to model potential interactions between the proteins (28). We used this method because there appears to be little protein sequence conservation of key Hook, Pad, or LolA residues identified in the *E. coli* structure among evolutionarily diverse bacteria (**Fig. S1-S2**).

Prediction of the *E. coli* LolC:LolA complex very closely matched the crystalized interaction captured between purified LolA and the LolC periplasmic domain (**Fig. 2A**). It is important to note that the AlphaFold2 algorithm was not trained on the LolC:LolA crystal structure which was released at a later date (25). Modeling predictions found practically the same contacts between the *E. coli* Pad and LolA (**Fig. S1-S2**). Examining the predictions of LolC/LolF:LolA interactions among diverse species, we observed that the insertion of the LolC/LolF Hook into the hydrophobic cavity of LolA is predicted to be a universal feature, despite the relatively low conservation of Hook amino acid sequences (**Fig. 2B** and **Fig. S1-S2**).

**Figure 2.**
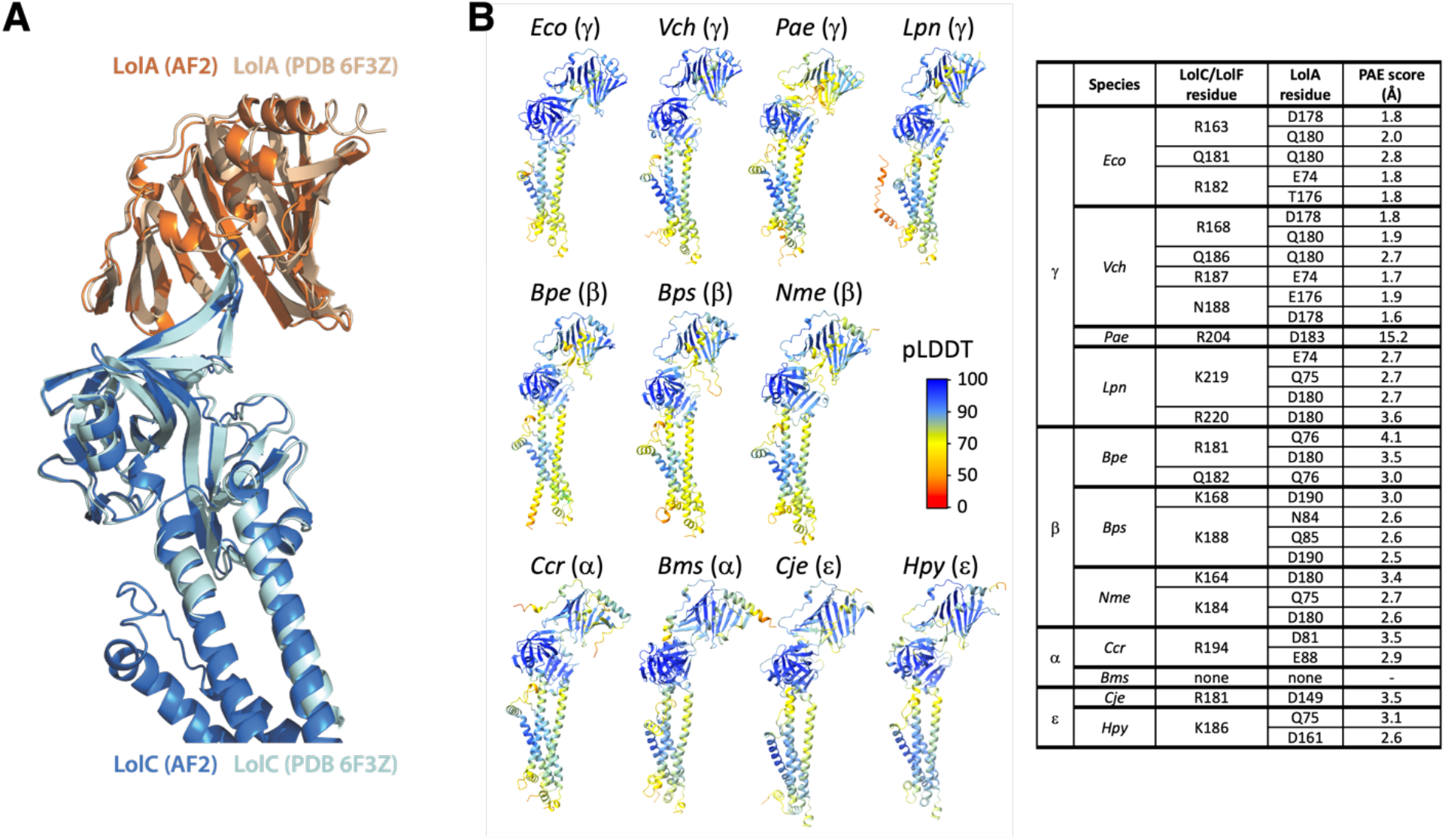
Predicted interactions between LolC/LolF and LolA among diverse bacteria. **(A)** AlphaFold2 Multimer prediction of *E. coli* LolA interacting with LolC aligned with the crystal structure of LolA co-crystalized with LolC periplasmic domain (PDB 6F3Z). **(B)** Aligned AlphaFold2 Multimer predictions of LolA proteins (top) interacting with their LolC/LolF partners (bottom). Insertion of the Hook sequence into the LolA cavity occurs in each prediction. Colors indicate pLDDT quality scoring for intrachain models. Table summarizes polar contacts between Pad regions and corresponding LolA proteins. The Predicted Aligned Error (PAE) score is a confidence prediction for an interaction between the described residue in the LolC Pad and the given LolA residue; PAE scoring ranges from 0-35 Å with scores <5Å having high confidence. LolA and LolC/F sequences are from *Escherichia coli* MG1655 (*Eco*), *Vibrio cholerae* O1 El Tor N16961 (*Vch*), *Pseudomonas aeruginosa* PAO1 (*Pae*), *Legionella pneumophila* subsp. pneumophila Philadelphia 1 (*Lpn*), *Bordetella pertussis* Tohama I (*Bpe*), *Burkholderia pseudomallei* K96243 (*Bps*), *Neisseria meningitidis* MC58 (*Nme*), *Caulobacter vibrioides* CB15 (*Ccr*), *Brucella suis* 1330 (Bms), *Campylobacter jejuni* subsp. jejuni NCTC 11168 (*Cje*), *Helicobacter pylori* 26695 (*Hpy*). Brackets denote Proteobacterial phyla of the organism. LolC producing bacteria are *Eco*, *Vch*, *Pae*. LolF producing bacteria are *Lpn*, *Bpe*, *Bps*, *Nme*, *Ccr*, *Bms*, *Cje*, *Hpy*.

Examining diverse predicted LolC/F Pad interactions with LolA, we saw differences compared to the interaction defined by crystallography of the *E. coli* components. Whereas the *E. coli* LolC Pad makes extensive polar contacts with C-terminal residues of LolA (aa 176-180) (**Fig. 2B**), the predicted models suggest that other species may rely on this LolA region significantly less or, in the case of α-Proteobacteria, not at all (**Fig. 2B**). In general, the models also predict fewer Pad residues participating in LolA interaction in distant relatives of *E. coli*. Analysis of the predictions suggests that the modes and extent of LolC Pad interaction with LolA may differ in other bacteria, perhaps suggesting a reduced reliance on the Pad in many species.

### The LolC Hook is essential for lipoprotein trafficking but is robust to mutation

We generated a series of mutations throughout the LolC periplasmic loop to assess their i*n vivo* effects; these mutations included those studied *in vitro* by Kaplan *et al*. (25) and others that we introduced in residues with outward-facing side chains. Given that LolC is essential, we expected that severely deleterious mutations would be lethal, while modestly deleterious mutations would impact growth. Since lipoprotein trafficking is needed for several OM assembly pathways, we expected mutations that impaired trafficking would disrupt OM integrity and cause sensitivity to large scaffold antibiotics that cannot normally penetrate the OM (e.g., vancomycin, rifampicin, or novobiocin). Based on the crystal structure of Hook:LolA, we mutated sites in the Hook that might disrupt Hook:LolA interaction (summarized in **Fig. 1B**). Mutations were introduced into *lolC* encoded within a *lolCDE* operon expressed from an arabinose-inducible promoter in the pBAD18 plasmid. In all, we generated 13 plasmid encoded LolC Hook variants and introduced each into wildtype *E. coli*.

To examine the *in vivo* effects of the *lolC* mutations, we first performed complementation tests with each of the LolC variants. Briefly, we generated P1*vir* phage that co-transduced Δ*lolCDE*::*cam* together with a nearby, neutral genetic marker (Δ*ycfR*::*kan*). After first selecting for the neutral marker (kanamycin-resistant transductants), we then screened for chloramphenicol-resistance to determine the frequency of Δ*lolCDE*::*cam* co-transduction. That is, we measured the genetic linkage between the two markers. Doubly-resistant transductants had recombined Δ*lolCDE*::*cam* to replace the native *lolCDE* operon and we interpreted such cells as tolerating deletion of native *lolCDE* because the plasmid-encoded *lolC* variant was able to support viability. As expected, control cells carrying plasmids encoding wildtype *lolCDE* readily tolerated inactivation of the native *lolCDE* locus, with ~79% of Δ*ycfR*::*kan* recombinants also recombining the Δ*lolCDE*::*cam* deletion-insertion allele (**Fig. 3A**).

**Figure 3.**
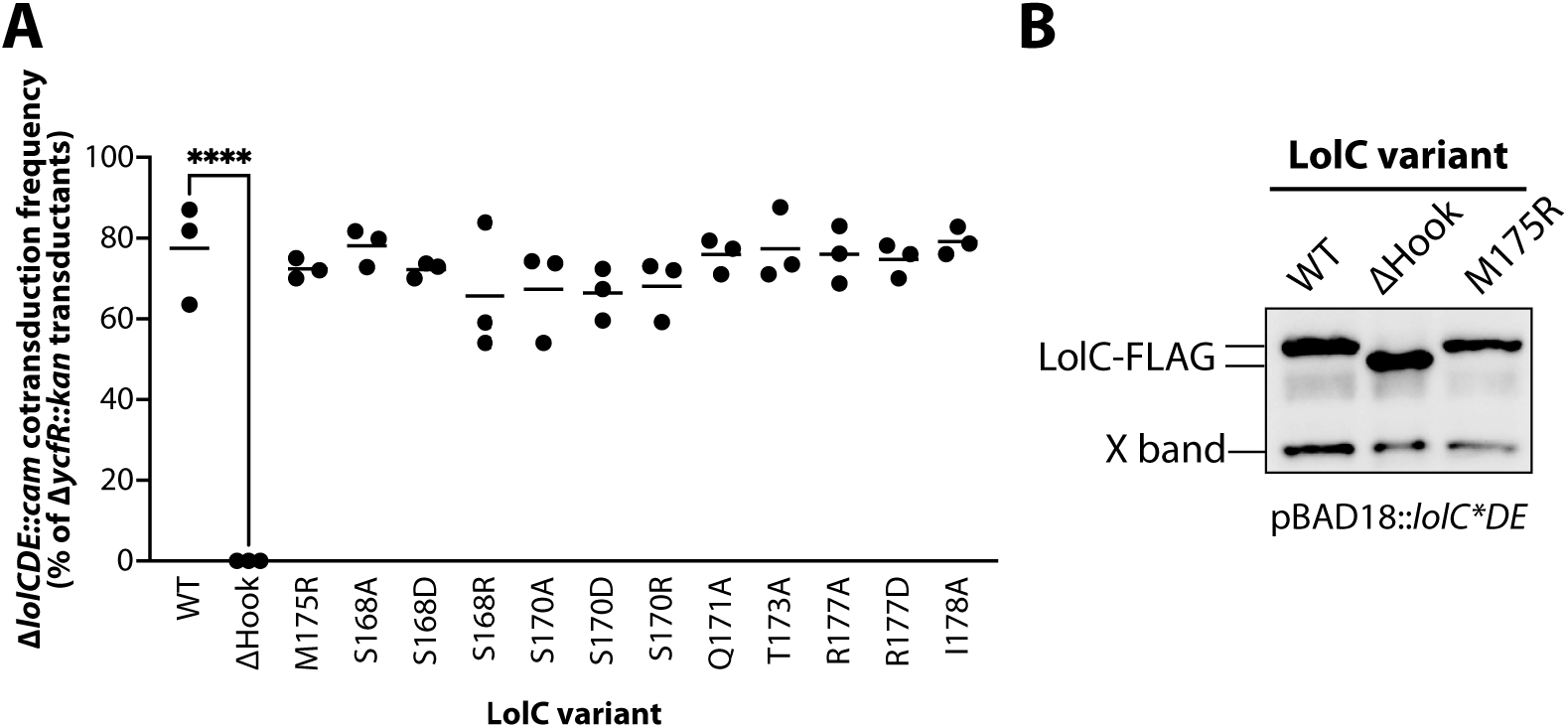
The Hook is essential for LolC activity. **(A)** Genetic linkage between Δ*ycfR*::*kan* and Δ*lolCDE*::*cam* when introduced into wildtype *E. coli* carrying pBAD18::*lolCDE* plasmids encoding the denoted *lolC* mutations. Three independent transductions were measured and 100 Kan^R^ transductants from each experiment were tests for Cam^R^ (each data point represents n=100); statistically significant results were found by one-way ANOVA (**** indicates *P<*0.0001). **(B)** Western immunoblotting with anti-FLAG antibodies of cell lysates from strains producing FLAG-tagged LolC proteins from pBAD18::FLAG-*lolCDE* plasmids. X-band is an *E. coli* protein that cross-reacts with anti-FLAG.

We found that production of LolC(ΔHook), which lacks residues 175-187, was unable to complement the loss of chromosomal *lolC*. Genetic linkage was severely disrupted; indeed, we did not recover any viable Δ*lolCDE*::*cam* transductants (**Fig. 3A**). One possible cause for this failed complementation may have been poor expression or stability of LolC(ΔHook) in *E. coli*. We constructed N-terminal FLAG-tagged LolC derivatives in pBAD18::*lolCDE* to assess protein levels by immunoblotting with anti-FLAG antibodies. We found that LolC(ΔHook) is expressed to levels that are comparable to LolC(wt) (**Fig. 3B**). Hence, we concluded that ΔHook reduces LolC function, impeding lipoprotein trafficking to an extent that cells are no longer viable.

Inefficient lipoprotein trafficking is known to cause toxic mislocalization of the highly abundant OM-targeted lipoprotein Lpp (19, 29). Under normal trafficking conditions, Lpp is efficiently trafficked to the OM and forms covalent attachments to the peptidoglycan cell wall via a C-terminal K58 residue (30, 31). When trafficking is inefficient, Lpp accumulates in the IM and forms cell wall attachments from this membrane, a reaction that is lethally toxic (19). Such toxicity can be abated by deleting the K58 residue (*lpp*(ΔK58)) to produce a highly abundant, though detoxified, OM-targeted Lpp variant (19). Alternatively, removing Lpp entirely (Δ*lpp*) can both prevent toxicity and substantially reduce the number of OM-target lipoproteins that require trafficking, since Lpp is the most abundant protein in the cell (32). We reasoned that LolC(ΔHook) failed to complement *lolC* either because the Hook is fundamentally essential for trafficking or because deletion of the Hook simply reduced trafficking efficiency to a level where Lpp mislocalization killed the cell. We tested that latter hypothesis by examining whether production of LolC(ΔHook) could complement loss of native *lolC* in both *lpp*(ΔK58) or Δ*lpp* backgrounds. Just as in wildtype (*lpp*^+^) *E. coli*, we failed to recover any Δ*lolCDE*::*cam* transductants (**Fig. S3**). This finding implied that the Hook is fundamentally essential, even when Lpp toxicity is abated and when there is reduced load on lipoprotein trafficking.

Given the *in vivo* importance of the Hook, we were surprised that none of the single substitution variants in the Hook displayed any defect in complementing loss of *lolC* that we could detect through genetic linkage (**Fig. 3A**). Similarly, we did not detect severe defects in growth (**Fig. 4A**) or OM permeability when *lolC* mutants were produced in haploid from plasmids (**Fig. 4B**). Most remarkable was LolC(M175R): when this substitution was present in a purified LolC periplasmic loop, it was found to completely abolish the loop’s ability to interact with purified LolA *in vitro* (25). Accordingly, we expected that the M175R substitution would similarly abolish LolA recruitment *in vivo* and that it would therefore be a lethal mutation. Instead, *E. coli* producing plasmid encoded LolC(M175R) were viable and only exhibited a modest growth defect (**Fig. 4A**) and appeared to produce a robust OM antibiotic barrier (**Fig. 4B**).

**Figure 4.**
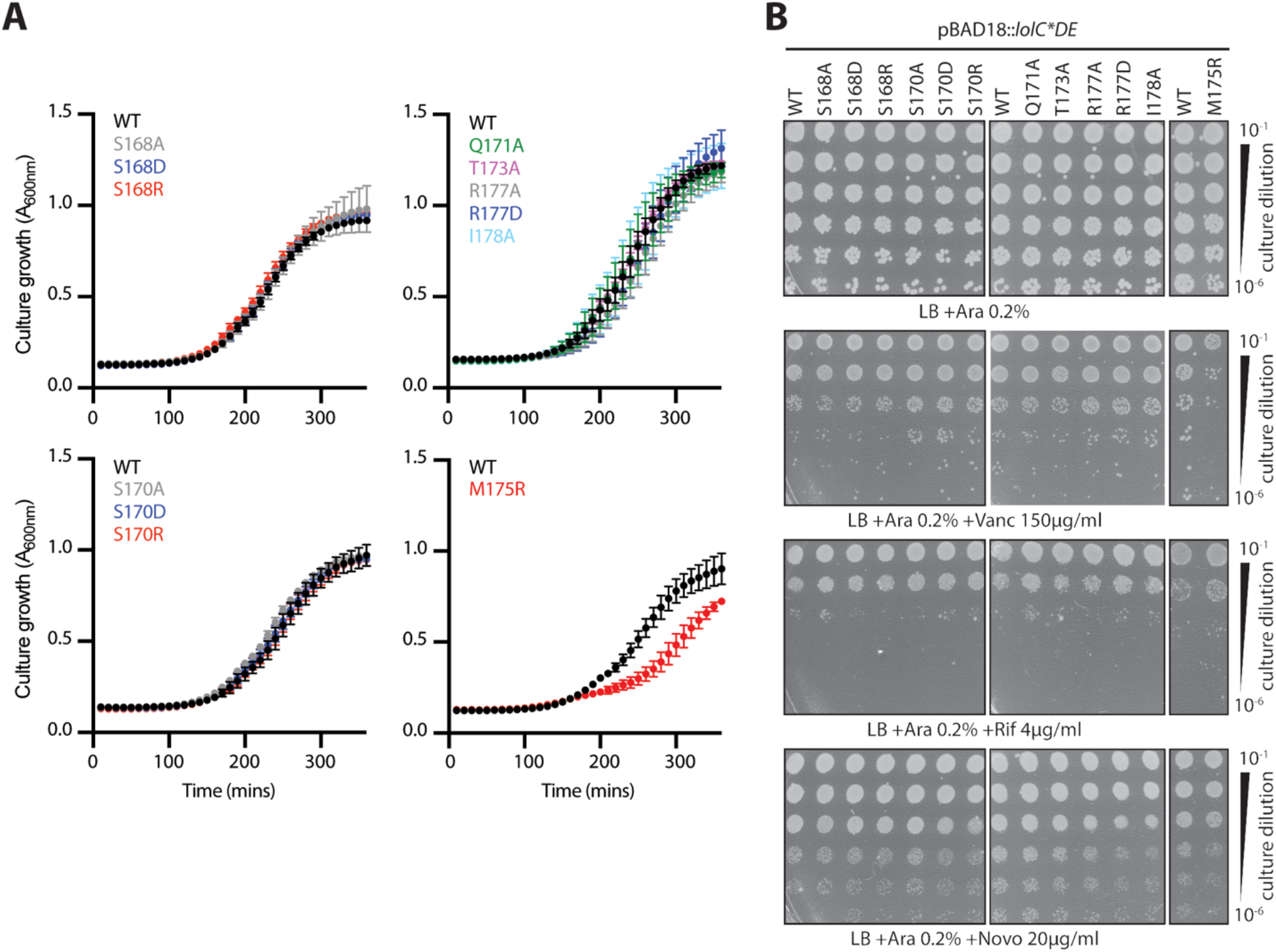
Individual Hook mutations have minimal impact on growth or OM permeability to antibiotics. **(A)** *E. coli* Δ*lolCDE*::*cam* strains carrying pBAD18::*lolCDE* plasmids with *lolC* mutations were grown in LB broth supplemented with 0.2% arabinose at 37°C. Data are average ± standard deviation (n=3). **(B)** Overnight cultures of *E. coli* Δ*lolCDE*::*cam* strains carrying pBAD18::*lolCDE* plasmids with *lolC* mutations were 10-fold serially diluted, spotted onto indicated agar plates, and incubated overnight at 37°C.

Since we had performed our testing with multi-copy plasmid encoded *lolC* variants produced from a heterologous promoter, we considered the possibility that these experimental conditions may mask any defects caused by LolC(M175R). Hence, we sought to introduce the M175R mutation into the native *lolC* locus to study this variant at physiological levels. Given biochemical evidence that an M175R abolishes LolA binding *in vitro*, we were wary that the M175R substitution might be lethal and hence not tolerated at the native *lolC* locus. Strikingly, we found that the mutation could readily be introduced into the *lolC* gene. Clearly, M175R does not abolish essential LolA recruitment by LolC. In fact, the chromosomal *lolC*(M175R) strain displayed only a minor difference in growth compared to a wildtype *lolC*^+^ strain (**Fig. 5A**) and, remarkably, exhibited no OM permeability defects to antibiotics (**Fig. 5B**).

**Figure 5.**
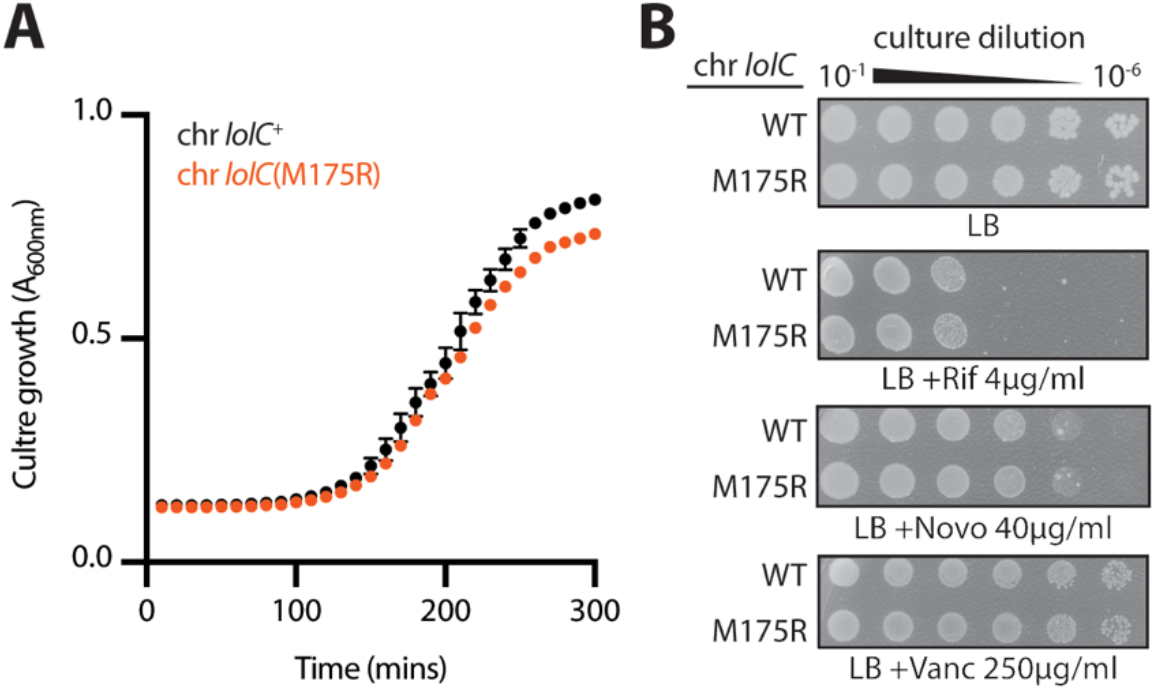
The M175R mutation is well-tolerated when introduced into the chromosomal *lolC* locus. **(A)** *E. coli* strains encoding either wildtype *lolC*^+^ or mutated *lolC*(M175R) at the native chromosome were grown in LB broth at 37°C. Data are average ± standard deviation (n=3). **(B)** Overnight cultures of *E. coli* strains encoding either wildtype or M175R chromosomal *lolC* alleles were 10-fold serially diluted, spotted onto indicated agar plates, and incubated overnight at 37°C.

As the chromosomal *lolC*(M175R) mutation was unexpectedly well tolerated, we constructed multiple substitution mutations in the Hook to further assess its importance for essential lipoprotein trafficking. First, we constructed a triple mutant of T173A M175R I178A; these substitutions were individually the most highly deficient for LolA binding *in vitro* (25). Second, we also constructed a complete alanine replacement of the Hook residues, where each residue in LolC(168-178) was substituted to alanine (with the exception of P174 at the apex turn of the Hook); we refer to this multi-alanine mutant as LolC(Hook-Ala). Producing FLAG-tagged LolC(T173A M175R I178A) or LolC(Hook-Ala) variants from pBAD18, we found that both LolC protein variants were well expressed in *E. coli*, with levels comparable to wildtype LolC (**Fig. 6A**). Using genetic linkage analysis to test if either LolC variant, when expressed from multicopy pBAD18 plasmid, could complement the loss of native *lolC*, we found that LolC(Hook-Ala) was unable to complement inactivation of native *lolC* (**Fig. 6B**). Indeed, LolC(Hook-Ala) failed to complement even in a Δ*lpp* strain where the number of lipoproteins that require trafficking is highly reduced (**Fig. S4**). Clearly, the complete replacement of Hook residues with alanines severely reduced function, to a level that is insufficient to sustain *E. coli* viability. The chemical identify of the Hook residues must therefore be important for LolC function. Surprisingly, we found that LolC(T173A M175R I178A) expressed from pBAD18 appeared to readily complement loss of native *lolC* even in *lpp*^+^ *E. coli* (Fig. 6B). However, we found that the Δ*lolCDE*::*cam* transductants grew very poorly despite expressing LolC(T173A M175R I178A) from a multicopy plasmid (**Fig. 6C**) and produced heterogenous colony sizes indicating poor viability (**Fig. 6D**). Indeed, we observed that the T173A M175R I178A mutation could not be introduced into the native *lolC* locus of *E. coli*, not even in Δ*lpp* cells. Collectively, our analyses of LolC Hook mutations underscore the essentiality of the Hook for lipoprotein trafficking. While the Hook is remarkably robust *in vivo* to single substitutions, the Hook is clearly essential and multiple substitutions in the Hook can cause severe a loss of function.

**Figure 6.**
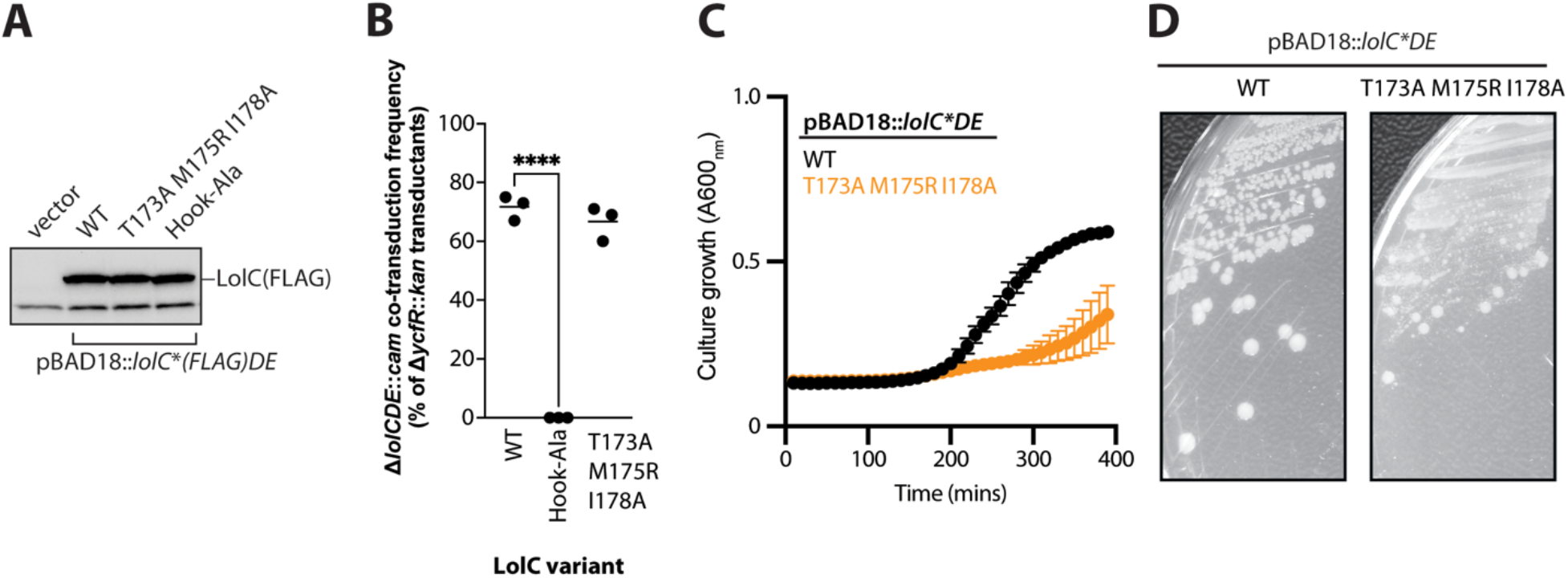
Multiple substitutions can inactivate the LolC Hook and impair viability. **(A)** Immunoblotting with anti-FLAG antibodies of cell lysates from strains producing FLAG-tagged LolC proteins from pBAD18::*lolC(FLAG)DE* plasmids. The lower band is an *E. coli* protein that cross-reacts with anti-FLAG. **(B)** Genetic linkage between Δ*ycfR*::*kan* and Δ*lolCDE*::*cam* when introduced into wildtype *E. coli* carrying pBAD18::*lolCDE* plasmids encoding the denoted *lolC* mutations. Three independent transductions were measured and 100 Kan^R^ transductants from each experiment were tests for Cam^R^ (each data point represents n=100); statistically significant results were found by one-way ANOVA (**** indicates *P<*0.0001). **(C)** *E. coli* Δ*lolCDE*::*cam* strains producing either wildtype *lolC*^+^ or mutated *lolC*(T173A M175R I178A) from pBAD18::*lolCDE* were grown in LB broth supplemented with 0.2% arabinose and incubated at 37°C. Data are average ± standard deviation (n=3). **(D)** *E. coli* Δ*lolCDE*::*cam* strains producing either wildtype *lolC*^+^ or mutated *lolC*(T173A M175R I178A) from pBAD18::*lolCDE* were streaked onto LB agar plates supplemented with 0.2% arabinose and incubated at 37°C. Highly heterogenous colony morphology is observed in the mutant strain.

### *In vivo* effects of mutations in the LolC Pad

The *E. coli* LolC Pad consists of three residues: R163 and adjacent residues Q181 and R182 (25). Biochemical experiments using the purified periplasmic domain of LolC and purified LolA found that alanine substitutions at any of these residues impaired LolA binding: R163A was a nonbinder, R182A was severely impaired, and Q181A was modestly impaired for LolA binding (25). We sought to examine the *in vivo* contribution of the Pad residues to lipoprotein trafficking. We mutated pBAD18::*lolCDE* to generate single alanine substitutions (R163A, Q181A, R182A), double alanine substitutions (R163A Q181A, R163A R182A), and a complete alanine replacement of all the Pad residues (R163A Q181A R182A). When FLAG-tagged, all these plasmid-encoded variants were produced at levels comparable to LolC(wt) (**Fig. 7A**).

**Figure 7.**
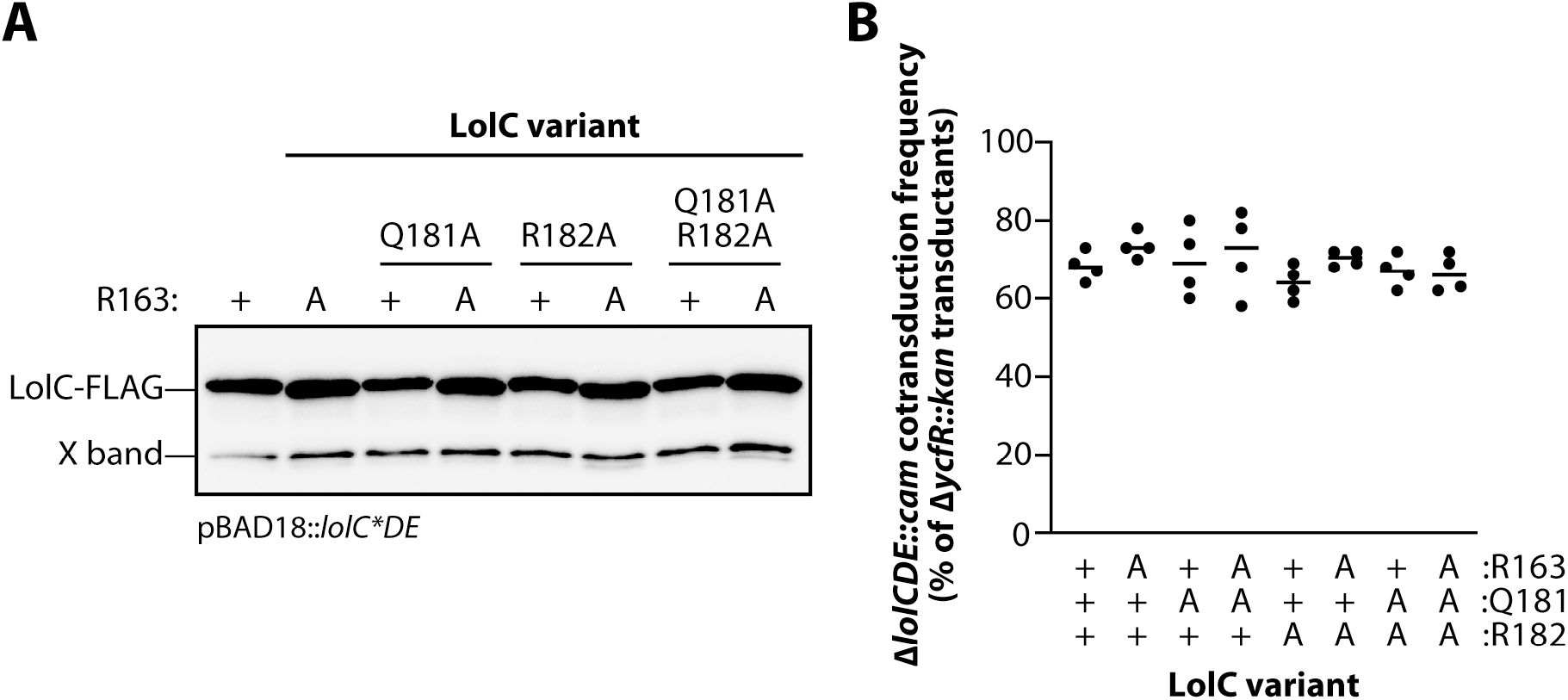
LolC Pad mutations expressed from plasmids are well tolerated in *E. coli*. **(A)** Western immunoblotting with anti-FLAG antibodies of cell lysates producing FLAG-tagged LolC proteins from pBAD18::FLAG-*lolCDE* plasmids. X-band is an *E. coli* protein that cross-reacts with anti-FLAG. **(B)** Genetic linkage between Δ*ycfR*::*kan* and Δ*lolCDE*::*cam* when introduced into wildtype *E. coli* carrying pBAD18::*lolCDE* plasmids encoding the denoted *lolC* mutations. Four independent transductions were measured and 100 Kan^R^ transductants from each experiment were tests for Cam^R^ (each data point represents n=100); one-way ANOVA detected no statistically significant differences.

First, we assessed the ability of plasmid-encoded Pad variants to complement inactivation of the native *lolC*. Surprisingly, we found that each of the variants complemented *lolC*, readily allowing for deletion of the native gene (**Fig. 7B**). This remarkable finding implied that the Pad was not required for any essential step in lipoprotein trafficking. However, we noticed that some mutants and, most notably, the complete replacement of the Pad residues with alanines (R163A Q181A R182A) caused a minor decrease in growth rate (**Fig. 8A**). Moreover, the combination mutations led to an impaired OM antibiotic barrier, leading to modest vancomycin sensitivity in a double mutant of R163A and R182A residues and in the triple Pad substitution mutant that additionally mutates Q181A (**Fig. 8B**). While modest, these phenotypes prompted us to re-examine the Pad residues at physiological levels.

**Figure 8.**
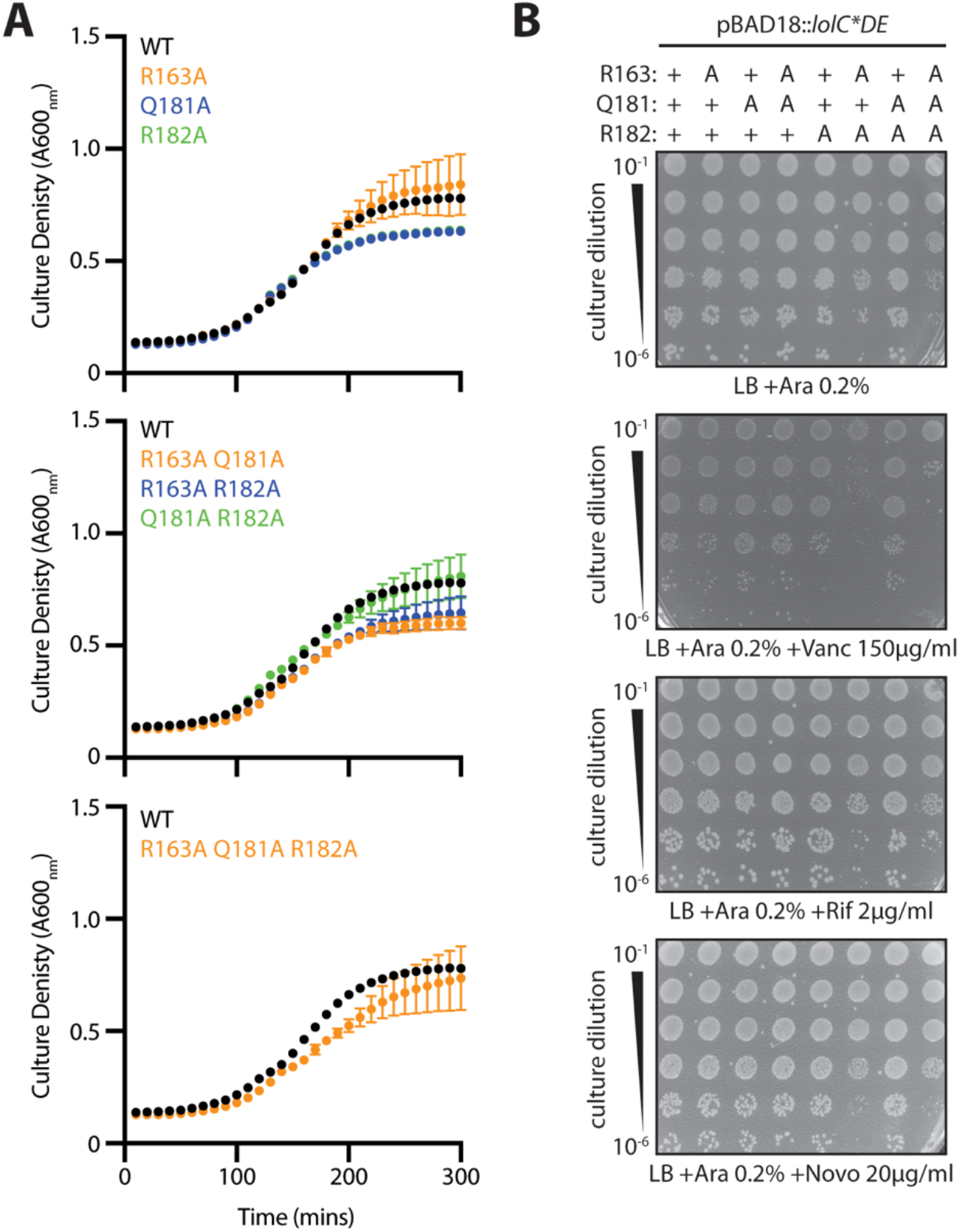
Pad mutations have minimal impact on growth or OM permeability to antibiotics. **(A)** *E. coli* Δ*lolCDE*::*cam* strains carrying pBAD18::*lolCDE* plasmids with *lolC* mutations were grown in LB broth supplemented with 0.2% arabinose at 37°C. Data are average ± standard deviation (n=3). **(B)** Overnight cultures of *E. coli* Δ*lolCDE*::*cam* strains carrying pBAD18::*lolCDE* plasmids with *lolC* mutations were 10-fold serially diluted, spotted onto agar plates, and incubated overnight at 37°C.

We first attempted to construct chromosomal *lolC* alleles encoding either a single R163A, a double Q181A R182A, or a triple R163A Q181A R182A substitution at the native locus. We were able to successfully construct *lolC*(R163A) and *lolC*(Q181A R182A) in a wildtype (*lpp*^+^) *E. coli* background, although *lolC*(R163A) caused a growth defect relative to wildtype cells (**Fig. 9A**). However, we repeatedly failed to recover any *lolC*(R163A Q181A R182A) mutants in *lpp*^+^ cells. Indeed, we could only construct the triple substitution mutant if the strain was complemented *in trans* with a wildtype *lolC* encoded on pBAD18::*lolCDE* (**Fig. 9B**). Hence, our data demonstrate that entirely inactivating the Pad with the triple substitution (R163A Q181A R182A) leads to a LolC variant that is insufficient to support cell viability in Lpp-producing *E. coli*.

**Figure 9.**
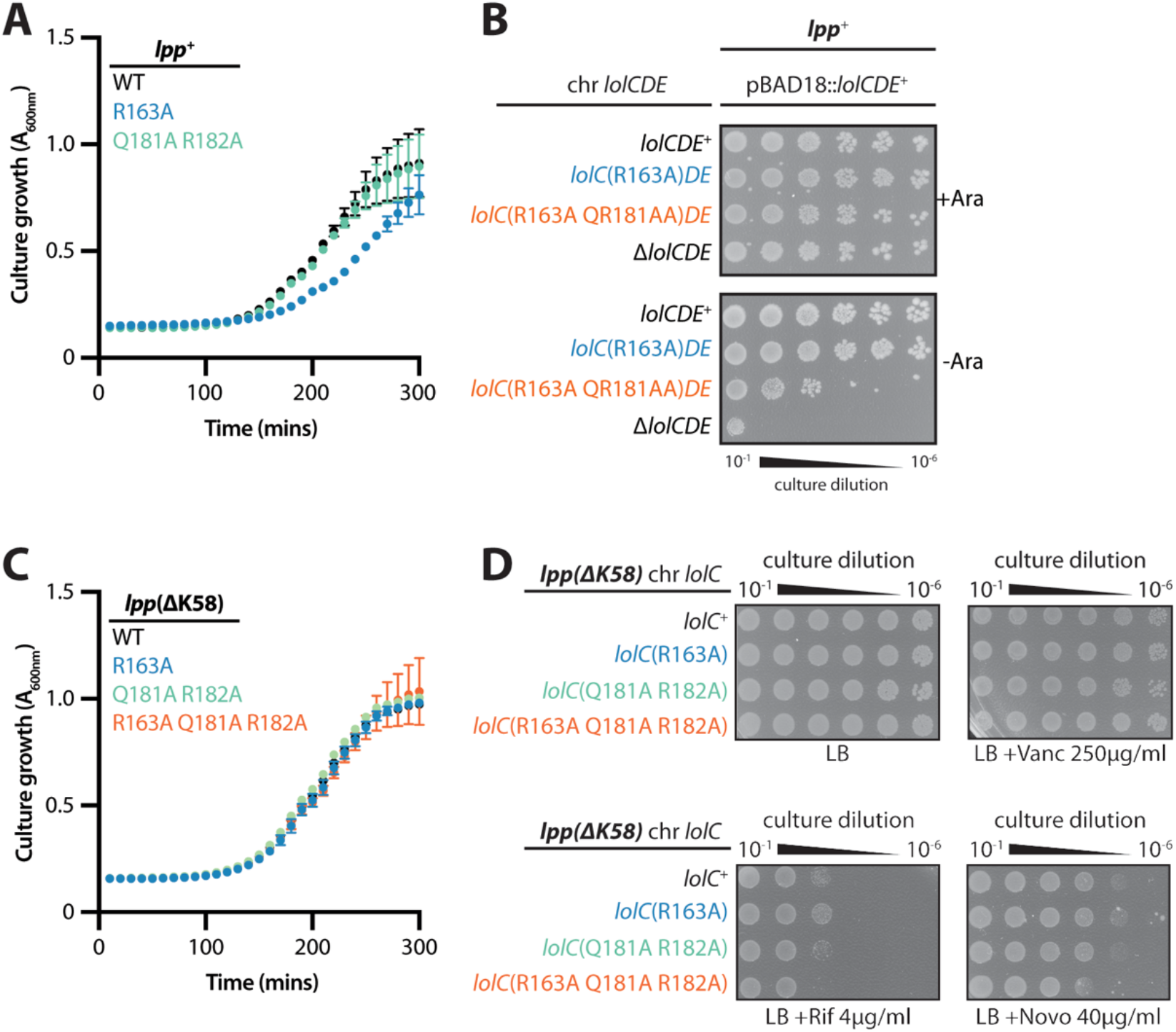
The LolC Pad is non-essential, but its inactivation causes Lpp-dependent lethality. **(A)** *E. coli* strains encoding *lolC* mutations in the native locus were grown in LB broth at 37°C. Data are average ± standard deviation (n=3). **(B)** Overnight cultures of *E. coli* encoding *lolC* mutations in the native locus were 10-fold serially diluted, spotted onto agar plates, and incubated overnight at 37°C. A chromosomal triple alanine Pad substitution *lolC* allele failed to support viability of *lpp*^+^ *E. coli*. **(C)** *E. coli lpp*(ΔK58) strains encoding *lolC* mutations in the native locus were grown in LB broth at 37°C. Data are average ± standard deviation (n=3). **(D)** Overnight cultures of *E. coli lpp*(ΔK58) strains encoding *lolC* mutations in the native locus were 10-fold serially diluted, spotted onto indicated agar plates, and incubated overnight at 37°C.

We reasoned that the *lolC*(R163A Q181A R182A) Pad triple mutant failed to support viability in *lpp*+ *E. coli* either: (i) because the Pad is fundamentally essential for a step in lipoprotein trafficking; or (ii) because the mutating the Pad caused less efficient lipoprotein trafficking that triggered lethal toxicity due to Lpp mislocalization. We sought to construct a *lolC*(R163A Q181A R182A) in an *E. coli lpp*(ΔK58) strain which still produces the highly abundant OM-targeted Lpp protein but in a form that cannot cross-link to the cell wall. To our surprise, we could readily introduce *lolC*(R163A Q181A R182A) and other Pad mutations into the native *lolC* locus in a *lpp*(ΔK58) strain. Moreover, none of the Pad mutants displayed any defect in growth in *lpp*(ΔK58) *E. coli* (**Fig. 9C**), suggesting that lipoprotein trafficking was not severely impacted by the loss of native Pad residues. Likewise, we detected no significant defect in OM permeability against antibiotics in the chromosomal Pad mutants in *E. coli lpp*(ΔK58) (**Fig. 9D**). We earlier detected modest vancomycin sensitivity with some plasmid-encoded Pad mutants (**Fig. 8B**); since this defect was not detected with corresponding chromosomal *lolC* mutants, we think that non-physiological LolCDE overproduction in the plasmid system contributed to that phenotype.

### Loss of the LolC Pad only weakly impairs lipoprotein trafficking to the OM

The fact that we were unable to observe significant defects in viability, growth, or OM permeability in chromosomal triple Pad mutants (in Lpp(ΔK58) *E. coli*) was truly surprising given that even just one mutation in the LolC Pad had previously been shown to completely abolish *in vitro* binding of LolA ((25). To further investigate this unexpected finding, we first examined cellular levels of the essential OM β-barrel LptD that is essential for LPS transport. LptD undergoes a complex assembly, requiring two initial disulfide bonds, folding into the OM, and then re-arrangement of the disulfide bonds. Successful assembly of the mature, correctly oxidized, LPS transport-competent LptD (termed LptDox) requires the activities of several OM lipoproteins: the BAM complex lipoproteins are needed for folding into the OM (33); the LptE lipoprotein is needed for correct LptD folding and disulfide rearrangements (33); and the novel OM lipoprotein YifL/LptM is also needed for LptD oxidation (34). Hence, we expected that levels of LptDox should be highly sensitive to defects in OM lipoproteins trafficking which would impair the delivery of multiple lipoprotein factors needed for LptD maturation. We assayed LptDox levels in whole cells by performing non-reducing SDS-PAGE and immunoblotting. Notably, levels of LptDox in a chromosomal triple Pad mutant *lolC*(R163A Q181A R182A) *lpp*(ΔK58) strain were indistinguishable from those in either an isogenic *lolC*^+^ *lpp*(ΔK58) strain or true a WT strain (*lolC*^+^ *lpp*^+^) (**Fig. 10A**). These data suggested that the triple Pad mutant did not cause severe defects in OM lipoprotein trafficking, since a process heavily dependent on OM lipoproteins (LptD maturation) appeared to be unaffected.

**Figure 10.**
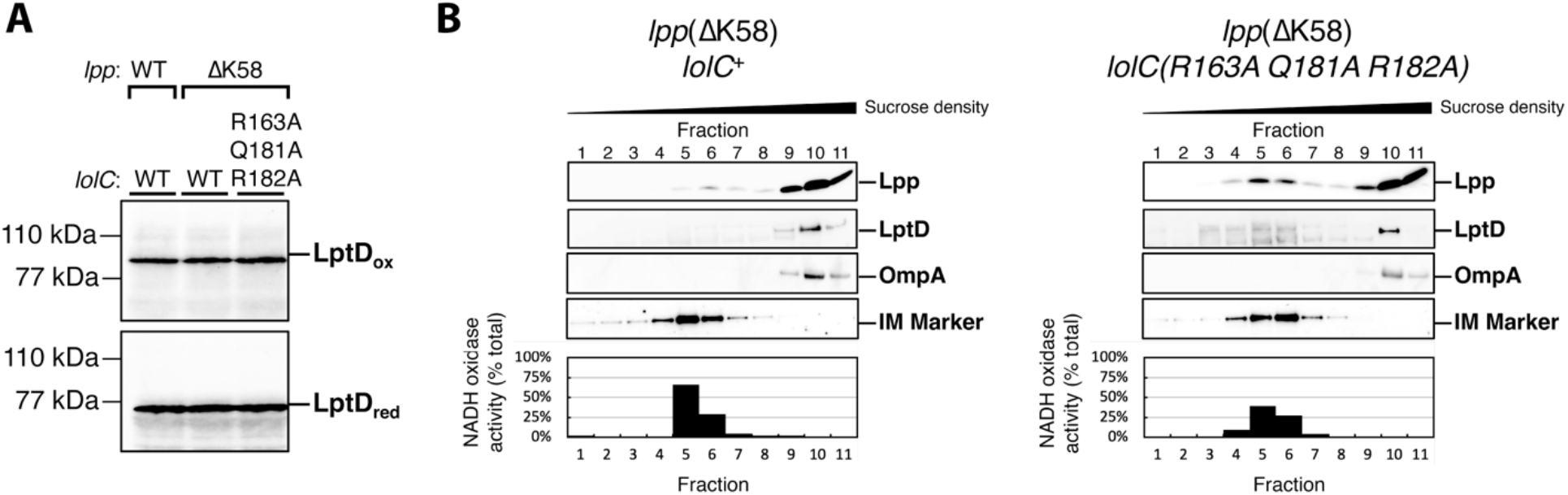
Inactivation of the LolC Pad causes only a minor defect in OM lipoprotein localization. **(A)** Levels of LptD β-barrel OM protein were measured from whole cells lysates by immunoblotting following either non-reducing SDS-PAGE (to detect LptDox) or reducing SDS-PAGE (to detect total LptD levels). **(B)** Inner and outer membranes from cell lysates were separated according to density using sucrose gradient centrifugation. Fractions of the gradient were collected and analyzed for composition. IM-containing fractions were identified by measuring NADH oxidase activity that only occurs in this membrane; the amount of total NADH activity per fraction is shown. An additional marker for the IM fraction was provided by immunoblotting with anti-LptD antisera and identifying the cross-reacting 55 kDa protein that was previously determined to be in the inner membrane (“IM Marker”) (59). OM-containing fractions were identified by immunoblotting against β-barrel OMPs LptD and OmpA. The localization of OM-targeted lipoprotein Lpp(ΔK58) was determined by immunoblotting fractions with anti-Lpp antisera. As expected, OM and IM fractions could be isolated from distinct fractions corresponding to high and low sucrose density, respectively.

To assess lipoprotein trafficking more directly, we next used sucrose density gradient fractionation to separate the inner and outer membranes and then examined the levels of the Lpp(ΔK58) lipoprotein in both membranes of each strain (**Fig 10B**). We found that levels of Lpp(ΔK58) in the OM of the *lolC*(R163A Q181A R182A) Pad mutant were similar to the *lolC*^+^ control strain. However, a small but notable increase in inner membrane accumulation of Lpp(ΔK58) could be detected in the Pad mutant compared to *lolC*^+^. Clearly, the Pad mutant is not as efficient in trafficking Lpp(ΔK58) to the OM as wild-type LolC; yet, the amount of mislocalized Lpp(ΔK58) is only a small fraction of this highly abundant lipoprotein which is mostly correctly trafficked to the OM. Since we found that production of wildtype Lpp is lethal in the Pad mutant, we infer that the small amount of inner membrane mislocalized Lpp(ΔK58) that we detected reflects the small amount of full-length Lpp that mislocalizes in the *lpp*+ Pad mutant strain and which is sufficient to be lethally toxic.

Collectively, our data demonstrate that replacement of the LolC Pad causes only a very minor defect in lipoprotein trafficking, as judged by the efficient trafficking of highly abundant Lpp(ΔK58) to the OM. That finding is consistent with no discernible defects in LptDox assembly, no appreciable defects in either viability or growth rate, and no OM permeability defects. However, our findings demonstrate the LolC Pad residues do not play a role in any essential step in lipoprotein trafficking to the OM. Rather, the Pad residues act to modestly enhances the efficiency of trafficking. The reduced efficiency when the Pad is inactivated can be lethal in the context of Lpp, and wildtype *E. coli* clearly require the Pad. However, cells lacking the LolC Pad still exhibit highly efficient OM lipoprotein trafficking, with only a minor mislocalization defect being detected.

## DISCUSSION

OM assembly in Gram-negative bacteria relies extensively on the trafficking of essential lipoprotein components (10, 35, 36). As a result, the lipoprotein maturation and trafficking pathways have been a focus of novel antibiotic discovery (6). Several compounds have recently been described that target maturation steps in both *E. coli* and in *Acinetobacter baumannii* (37–40). Novel inhibitors of lipoprotein trafficking have also been reported that target LolCDE in *E. coli* (21–23). However, each of these compounds appeared to have limited utility against *E. coli* due to rapid acquisition of *lpp* null mutations that confer resistance. These mutations likely allow cells to bypass of the toxic consequences of Lpp mislocalization caused by inhibition of OM lipoprotein trafficking. However, while *lpp* null mutations are well-tolerated in laboratory conditions, there is strong evidence that *lpp* mutants are severely compromised in virulence models in *E. coli* and closely related, *lpp*-encoding bacteria (41–46). Hence, *lpp* null mutations would be unlikely to arise in clinically relevant niches. Moreover, only a few closely related species produce Lpp, so the concern over rapid *lpp* resistance does not apply to many important pathogens. The recent emergence of structural information about the LolCDE transporter and how it recruits LolA could help to further refine efforts at inhibiting this early trafficking step (25, 26, 47, 48). What has been missing is a clear understanding of how structural and biochemical insights are reflected *in vivo*. The findings in our study provide this crucial *in vivo* context and redefine the current model of LolC:LolA interaction.

Two regions of the large LolC periplasmic loop had been suggested to recruit LolA. In biochemical assays, mutations in either region abolished the interaction between LolC and LolA *in vitro*, implying that both regions are essential for the process (25). However, rather than equally contributing to the recruitment of LolA, our genetic analysis demonstrates a hierarchy of importance that distinguishes the Hook from the Pad. Our data show that the Hook is fundamentally essential for the cell; the Pad cannot compensate for its absence. On the other hand, the Pad is not essential for trafficking. While alanine replacement of all the Pad residues is not tolerated in *lpp*^+^, removing a single codon in *E. coli* (specifying K58 of Lpp) is sufficient to fully obviate the requirement for the native Pad residues. Importantly, even in *lpp*(ΔK58) cells, lipoprotein trafficking via LolA is essential for *E. coli* viability (19); hence, LolA recruitment by LolC remains an essential trafficking step in *lpp*(ΔK58). The fact that a chromosomal *lolC* triple alanine substitution mutation of the Pad residues is viable, exhibits only a minor defect in lipoprotein OM localization, shows no growth defects, and displays no OM permeability defects in *lpp*(ΔK58) underscores the non-essential contribution of the LolC Pad in lipoprotein trafficking.

Our modeling of diverse LolC/F:LolA recruitment interaction models are congruent with the idea that the Hook is the main player in LolA recruitment. All the predicted models place the Hook within the hydrophobic cavity of LolA where it makes extensive interactions. Conversely, predictions of Pad residue interactions with LolA vary. AlphaFold2 was remarkably good at predicting the crystalized *E. coli* Pad:LolA interaction. It is notable, therefore, that AlphaFold2 predicts fewer sites of interaction between diverse LolA proteins and their corresponding LolC/LolF Pad regions. While testing of these models remains ongoing, we hypothesize that the predictions reflect a less important (and perhaps ancillary) role for the Pad in recruiting LolA among diverse bacteria. Indeed, our *in vivo* data suggests that replacing all the Pad residues with alanines does not abolish LolC activity, but modestly reduces it to a level incapable of dealing with Lpp, a lipoprotein that is absent in all the modeled species except for *E. coli* and *V. cholerae*. Our finding that the LolC Pad is dispensable is also consistent with prior LolA C-terminal truncation studies where removal of the final five amino acids was well tolerated *in vivo* in wildtype *lpp*^+^ *E. coli* (49). Crucially, that LolA truncation removed the interacting residues for LolC(R163) which *in vitro* biochemical assays found to be essential for LolA recruitment (25). In fact, even a larger nine amino acid truncation that removed all the C-terminal LolA residues that interact with the LolC Pad was found to be viable in *E. coli* (at least when grown on agar) (49).

Our study underscores two considerations for genetic or biochemical study of LolCDE. First, our findings are another reminder for the need to carefully consider multicopy effects when studying LolCDE. Since we can readily detect plasmid expressed FLAG-LolC but have been unable to detect FLAG-LolC produced from the native locus (not shown), it is clear that the plasmid produces significantly more LolCDE than endogenous levels and this can obscure phenotypic defects. Indeed, multicopy overexpression of LolCDE was even previously shown to enable trafficking of non-native diacylated lipoproteins (50). Careful analysis is needed to pinpoint subtle deficiencies for subsequent investigations of *lolCDE* at native levels. Second, our findings are another indication that current biochemical assays of lipoprotein trafficking (reported by us and others) appear to have a low dynamic range and are highly sensitive to disruption (51, 52). The trafficking system *in vivo* is remarkably robust by comparison. This finding has implications for *in vitro* approaches to identify disruptors of LolC:LolA interaction, such as peptide screening.

LolC recruitment of LolA is an essential early step in lipoprotein trafficking. Coalescing structural, biochemical, and genetic advances now leads us to propose a new model for LolA recruitment. We propose that the LolC Hook recruits LolA with high affinity; the Hook makes extensive contacts within the hydrophobic LolA cavity. It appears that the Hook interaction is difficult to displace through individual mutational changes. All our Hook substitutions were well tolerated, even an M175R substitution (that abolished LolA recruitment *in vitro*) could readily be introduced into the native *lolC* gene. Meanwhile, we propose that the LolC Pad aids in LolA recruitment, perhaps by positioning the C-terminal LolA region so that it can efficiently receive incoming lipoprotein cargo. This contribution of the Pad would likely enhance trafficking efficiency. Ultimately, however, the role of the Pad is secondary to the primary role of the Hook and the Pad’s inactivation presents only modest impairment of OM lipoprotein trafficking.

## MATERIALS AND METHODS

### Bacterial strains, plasmids, and growth conditions

Strains and plasmids used in this study are listed in **Tables S1** and **S2**, respectively. Chromosomal mutant alleles were introduced by P1*vir* transduction. Null alleles were obtained from the Keio collection (53) and their Kan^R^ cassettes were cured using plasmid pCP20 (54). Δ*lolCDE*::*cam* was described previously (19). *zce*-*726*::Tn*10* was also described previously (55). Unless otherwise indicated, strains were routinely grown in Lennox broth (LB) or agar at 37°C. LB was supplemented with ampicillin (Amp; 25 g/ml), chloramphenicol (Cam; 20 g/ml), kanamycin (Kan; 25 g/ml), tetracycline (Tet; 25 g/ml), and L-arabinose (0.2% [wt/vol]) as required.

### Construction of plasmid encoded LolC variants

Plasmid pBAD18::*lolCDE* was subjected to PCR-based site-directed mutagenesis using the oligonucleotides listed in **Table S3** to generate each corresponding *lolC* mutation. Plasmids were Sanger sequenced across the *lolC* gene to confirm presence of the mutation. FLAG-tagged LolC variants were constructed by an in-frame insertion of sequence encoding an N-terminal FLAG epitope immediately after the ATG initiation codon. Using chromosomal alleles, we confirmed that N-terminal FLAG-tagging of LolC did not adversely impact growth or OM antibiotic permeability, relative to native untagged LolC (**Fig. S5**)

### Construction of chromosomal LolC variants

To generate double stranded DNA templates for introducing *lolC* mutations into the native site by recombination, we PCR amplified plasmid-encoded *lolC* variants with primers rec_p18lolCDE_fwd and rec_p18lolCDE_rev. The primers amplify the entire *lolCDE* operon and provide 120 bp and 107 bp of flanking 5’ and 3’ homology, respectively. Purified PCR products were used to transform a derivative of the recombinogenic strain DY378 as described previously (56). The DY378 derivative strain was constructed by first transforming it with pBAD18::*lolCDE* and then transducing it with Δ*lolCDE*::*cam*. Growth of the resulting strain is dependent on arabinose supplemented to media. The strain’s λRed recombinase was induced as described previously (56), electrocompetent cells were prepared and PCR DNA for recombination was electroporated into the cells. Transformants were selected by plating onto LB agar lacking any arabinose. Successful recombinants at the *lolCDE* locus lost arabinose-dependence and were Cam^S^. Recombinants were PCR screened with primers lolCDE_screen_F and lolCDE_int1_R and the products were Sanger sequenced to confirm presence of the intended mutation. *lolC* mutations were genetically linked to *zce-726*::Tn*10* to facilitate movement of alleles into other *E. coli* backgrounds by co-transduction. Chromosomal *lolC* mutant strains were confirmed by whole-genome sequencing.

Construction of chromosomal alleles was also performed with the same scheme in Δ*lpp*::*kan* derivatives of the recombinogenic strain.

### Genetic linkage analysis

To test the ability of pBAD18::*lolCDE* plasmid-encoded *lolC* variants to complement loss of the native *lolC* gene, *E. coli* strains carrying the plasmid were grown overnight in LB supplemented with Amp. 0.1 ml of saturated culture was transduced with P1*vir* phage generated from a donor encoding linked markers *zce-760*::Tn*10* Δ*lolCDE*::*cam*. Tet^R^ transductants were selected on LB agar (also supplemented with arabinose). 100 transductants were screened for Cam^R^ on LB agar (also supplemented with arabinose) to test for co-transduction of Δ*lolCDE*::*cam*. The frequency of co-transduction was measured for at least 3 independent biological replicates. Statically significant differences in co-transduction were detected using one-way ANOVA analysis.

### Growth analysis

Saturated overnight cultures were subcultured into fresh broth at a dilution of 1:1,000 and seeded (2 ml) into wells of a 24-well microtiter plate. Cultures were grown with shaking at 37°C in a BioTek H1 or BioTek Epoch2 multimode incubating plate reader. Growth was assessed from independent biological replicates. Standard deviation was calculated.

### Sucrose density gradient fractionation

Separation of inner and outer membranes was performed as described by (57) with minor modifications. For each strain, a 250-ml culture (inoculated from an overnight culture at a 1:100 dilution) was grown in LB broth until reaching an A600nm of 0.6–0.8. Cells were washed once with 25 ml of cold 10mM Tris-HCl, pH 8.0, and centrifuged at 10 000 x g for 10 min at 4°C. Cells were resuspended in 20 ml of 10 mM Tris-HCl, pH 8.0 containing 20% sucrose (w/w), Benzonase (EMD Millipore) and HALT protease inhibitor cocktail (Thermo Scientific), and then lysed by a single passage through a French Pressure Cell Press (Thermo Spectronic) at 8000 psi. Unbroken cells were removed by centrifugation at 10 000 x g for 10 min at 4°C. The cleared cell lysate was collected, and 5.5 ml was layered on top of a two-step sucrose gradient consisting of 5 ml of 40% sucrose solution layered on top of 1.5 ml of a 65% sucrose solution in a 14 x 89mm Ultra-Clear tube (Beckman Coulter). All sucrose (w/w) solutions were prepared in 10mM Tris-HCl, pH 8.0. Samples were centrifuged at 35 000 rpm for 18 h in a Beckman SW41 rotor in an OptimaXE-90 Ultra-centrifuge (Beckman Coulter). Eleven fractions were manually collected from each tube, starting from the lowest density i.e., from the top of the tube.

### NADH Oxidase Assay

The inner membrane enzyme, NADH oxidase, was used as a marker for IM material as previously described by (58) with minor modifications. Briefly, 2.5 *μ*L of each fraction from the sucrose density gradient was added to a 96-well black bottom plate and 97.5 *μ*L of 100 mM Tris HCl, pH 8.0, 120 *μ*M NADH (Sigma) and 5 mM dithiothreitol (Sigma) was added to each well. Changes in fluorescence (ex. = 340 nm and em. = 465 nm) were monitored over the course of 2 minutes. The activity of NADH oxidase for each fraction is represented as a % of total NADH oxidase activity in the gradient.

### Whole-cell lysate preparation and immunoblotting

Equivalent cell densities (normalized by OD 600nm) were pelleted by centrifugation (10,000 x g for 5 min). Pellets were resuspended in 2x Tris-Glycine SDS Sample Buffer with 4% beta-mercaptoethanol (B-ME). Lysates were incubated at 100°C for 5 min. Samples were resolved on 4-20% Tris-Glycine gels (Novex) at 200 V for 45 min. For immunoblotting of LPP, gradient fractions were prepared in Tricine Sample Buffer (Novex) and resolved on 16% Tricine gels (Novex). Protein samples resolved by polyacrylamide gel electrophoresis were transferred to 0.2 μm nitrocellulose membranes and probed with either anti-FLAG M2 monoclonal antibodies (Sigma-Aldrich), used at a dilution of 1:10,000), anti-Lpp (Silhavy Lab stock, used at a dilution of 1:400 000), or anti-IMP/LptD rabbit polyclonal anitsera (Silhavy Lab stock, used at a dilution of 1:5,000). The anti-IMP/LptD polyclonal antibody also reacts with a 55-kDa protein IM protein and OmpA 37 kDa (59). Membranes were subsequently probed with either goat anti-mouse-HRP (BioRad) or goat anti-rabbit-HRP secondary antibodies (EMD Millipore) as appropriate. Probed membranes were developed by incubating with Immobilon Classico Western HRP substrate (EMD Millipore). The resulting chemiluminescence was detected with a BioRad ChemiDoc MP.

### AlphaFold2 Multimer predictions

To obtain mature sequences of each LolA protein, the site of Signal Peptidase I cleavage was predicted with SignalP-6.0 (60). Mature LolA and complete LolC/F sequences from each species were used to predict potential interactions using a simplified version of the AlphaFold2 v2.3.2 multimer extension (28). The program exported the model with the highest confidence (pTM). A summary of pLDDT and PAE intra- and inter-chain prediction quality assessments are provided in Fig. 2B.

### Efficiency of plating assays

Efficiency of plating assays were used to determine the relative sensitivities of strains to vancomycin, rifampin and novobiocin. Assays were performed by preparing 10-fold serial dilutions of overnight cultures (standardized by A_600nm_) in 96-well microtiter plates before replica plating onto non-selective LB agar (supplemented with arabinose, as required) and selective antibiotic-containing agar medium. Plates were then incubated overnight at 37°C.

## ACKNOWLEDGEMENTS

This work was supported by grant R35 GM133509 (to M.G.) and fellowship F31 AI147589 (to K.M.L.). We thank all members of the Grabowicz lab for helpful discussions and comments on the manuscript.

## REFERENCES

1. Konovalova A, Kahne DE, Silhavy TJ. 2017. Outer Membrane Biogenesis. Annu Rev Microbiol 71:539–556.

2. Sun J, Rutherford ST, Silhavy TJ, Huang KC. 2022. Physical properties of the bacterial outer membrane. Nat Rev Microbiol 20:236–248.

3. Ruiz N, Wu T, Kahne D, Silhavy TJ. 2006. Probing the Barrier Function of the Outer Membrane with Chemical Conditionality. ACS Chem Biology 1:385–395.

4. Manrique PD, López CA, Gnanakaran S, Rybenkov VV, Zgurskaya HI. 2023. New understanding of multidrug efflux and permeation in antibiotic resistance, persistence, and heteroresistance. Ann New York Acad Sci 1519:46–62.

5. Collaborators AR. 2022. Global burden of bacterial antimicrobial resistance in 2019: a systematic analysis. The Lancet 399:629–655.

6. Lehman KM, Grabowicz M. 2019. Countering Gram-Negative Antibiotic Resistance: Recent Progress in Disrupting the Outer Membrane with Novel Therapeutics. Antibiotics (Basel, Switzerland) 8:163.

7. Theuretzbacher U, Blasco B, Duffey M, Piddock LJV. 2023. Unrealized targets in the discovery of antibiotics for Gram-negative bacterial infections. Nat Rev Drug Discov 10.1038/s41573-023-00791-6.

8. Okuda S, Sherman DJ, Silhavy TJ, Ruiz N, Kahne D. 2016. Lipopolysaccharide transport and assembly at the outer membrane: the PEZ model. Nat Rev Microbiol 14:337–345.

9. Doyle MT, Bernstein HD. 2022. Function of the Omp85 Superfamily of Outer Membrane Protein Assembly Factors and Polypeptide Transporters. Annu Rev Microbiol 76:259–279.

10. Grabowicz M. 2018. Lipoprotein Transport: Greasing the Machines of Outer Membrane Biogenesis. Bioessays 40:1700187.

11. Buddelmeijer N. 2015. The molecular mechanism of bacterial lipoprotein modification—How, when and why? FEMS Microbiol Rev 39:246–261.

12. Rayes JE, Rodríguez-Alonso R, Collet J-F. 2021. Lipoproteins in Gram-negative bacteria: new insights into their biogenesis, subcellular targeting and functional roles. Curr Opin Microbiol 61:25–34.

13. Grabowicz M. 2019. Lipoproteins and Their Trafficking to the Outer Membrane. Ecosal Plus 8:67–76.

14. Yakushi T, Masuda K, Narita S, Matsuyama S, Tokuda H. 2000. A new ABC transporter mediating the detachment of lipid-modified proteins from membranes. Nat Cell Biology 2:212–218.

15. Narita S, Tanaka K, Matsuyama S, Tokuda H. 2002. Disruption of lolCDE, Encoding an ATP-Binding Cassette Transporter, Is Lethal for Escherichia coli and Prevents Release of Lipoproteins from the Inner Membrane. J Bacteriol 184:1417–1422.

16. Ito Y, Kanamaru K, Taniguchi N, Miyamoto S, Tokuda H. 2006. A novel ligand bound ABC transporter, LolCDE, provides insights into the molecular mechanisms underlying membrane detachment of bacterial lipoproteins. Mol Microbiol 62:1064–1075.

17. Matsuyama S, Tajima T, Tokuda H. 1995. A novel periplasmic carrier protein involved in the sorting and transport of Escherichia coli lipoproteins destined for the outer membrane. The EMBO J 14:3365–3372.

18. Matsuyama S, Yokota N, Tokuda H. 1997. A novel outer membrane lipoprotein, LolB (HemM), involved in the LolA (p20)-dependent localization of lipoproteins to the outer membrane of Escherichia coli. The EMBO J 16:6947–6955.

19. Grabowicz M, Silhavy TJ. 2017. Redefining the essential trafficking pathway for outer membrane lipoproteins. Proc National Acad Sci 114:4769–4774.

20. Caro F, Place NM, Mekalanos JJ. 2019. Analysis of lipoprotein transport depletion in Vibrio cholerae using CRISPRi. Proc National Acad Sci 116:17013–17022.

21. Nayar AS, Dougherty TJ, Ferguson KE, Granger BA, McWilliams L, Stacey C, Leach LJ, Narita S, Tokuda H, Miller AA, Brown DG, McLeod SM. 2015. Novel Antibacterial Targets and Compounds Revealed by a High-Throughput Cell Wall Reporter Assay. J Bacteriol 197:1726–1734.

22. McLeod SM, Fleming PR, MacCormack K, McLaughlin RE, Whiteaker JD, Narita S, Mori M, Tokuda H, Miller AA. 2015. Small-Molecule Inhibitors of Gram-Negative Lipoprotein Trafficking Discovered by Phenotypic Screening. J Bacteriol 197:1075–1082.

23. Nickerson NN, Jao CC, Xu Y, Quinn J, Skippington E, Alexander MK, Miu A, Skelton N, Hankins JV, Lopez MS, Koth CM, Rutherford S, Nishiyama M. 2018. A Novel Inhibitor of the LolCDE ABC Transporter Essential for Lipoprotein Trafficking in Gram-Negative Bacteria. Antimicrob Agents Chemother 62:e02151–17.

24. Wilson A, Ruiz N. 2021. Transport of lipopolysaccharides and phospholipids to the outer membrane. Curr Opin Microbiol 60:51–57.

25. Kaplan E, Greene NP, Crow A, Koronakis V. 2018. Insights into bacterial lipoprotein trafficking from a structure of LolA bound to the LolC periplasmic domain. Proc National Acad Sci 115:E7389–E7397.

26. Tang X, Chang S, Zhang K, Luo Q, Zhang Z, Wang T, Qiao W, Wang C, Shen C, Zhang Z, Zhu X, Wei X, Dong C, Zhang X, Dong H. 2021. Structural basis for bacterial lipoprotein relocation by the transporter LolCDE. Nat Struct Mol Biol 28:347–355.

27. LoVullo ED, Wright LF, Isabella V, Huntley JF, Pavelka MS. 2015. Revisiting the Gram-Negative Lipoprotein Paradigm. J Bacteriol 197:1705–1715.

28. Jumper J, Evans R, Pritzel A, Green T, Figurnov M, Ronneberger O, Tunyasuvunakool K, Bates R, Žídek A, Potapenko A, Bridgland A, Meyer C, Kohl SAA, Ballard AJ, Cowie A, Romera-Paredes B, Nikolov S, Jain R, Adler J, Back T, Petersen S, Reiman D, Clancy E, Zielinski M, Steinegger M, Pacholska M, Berghammer T, Bodenstein S, Silver D, Vinyals O, Senior AW, Kavukcuoglu K, Kohli P, Hassabis D. 2021. Highly accurate protein structure prediction with AlphaFold. Nature 596:583–589.

29. Yakushi T, Tajima T, Matsuyama S, Tokuda H. 1997. Lethality of the covalent linkage between mislocalized major outer membrane lipoprotein and the peptidoglycan of Escherichia coli. J Bacteriol 179:2857–2862.

30. Braun V, Rehn K. 1969. Chemical Characterization, Spatial Distribution and Function of a Lipoprotein (Murein-Lipoprotein) of the E. coli Cell Wall. Eur J Biochem 10:426–438.

31. Asmar AT, Collet J-F. 2018. Lpp, the Braun lipoprotein, turns 50—major achievements and remaining issues. FEMS Microbiol Lett 365.

32. Li G-W, Burkhardt D, Gross C, Weissman JS. 2014. Quantifying Absolute Protein Synthesis Rates Reveals Principles Underlying Allocation of Cellular Resources. Cell 157:624–635.

33. Lee J, Xue M, Wzorek JS, Wu T, Grabowicz M, Gronenberg LS, Sutterlin HA, Davis RM, Ruiz N, Silhavy TJ, Kahne DE. 2016. Characterization of a stalled complex on the β-barrel assembly machine. Proc Natl Acad Sci 113:8717–8722.

34. Yang Y, Chen H, Corey RA, Morales V, Quentin Y, Froment C, Caumont-Sarcos A, Albenne C, Burlet-Schiltz O, Ranava D, Stansfeld PJ, Marcoux J, Ieva R. 2023. LptM promotes oxidative maturation of the lipopolysaccharide translocon by substrate binding mimicry. Nat Commun 14:6368.

35. Kim S, Malinverni JC, Sliz P, Silhavy TJ, Harrison SC, Kahne D. 2007. Structure and Function of an Essential Component of the Outer Membrane Protein Assembly Machine. Science 317:961–964.

36. Chng S-S, Ruiz N, Chimalakonda G, Silhavy TJ, Kahne D. 2010. Characterization of the two-protein complex in Escherichia coli responsible for lipopolysaccharide assembly at the outer membrane. Proc National Acad Sci 107:5363–5368.

37. Kitamura S, Owensby A, Wall D, Wolan DW. 2018. Lipoprotein Signal Peptidase Inhibitors with Antibiotic Properties Identified through Design of a Robust In Vitro HT Platform. Cell Chem Biology 25:301–308.e12.

38. Garland K, Pantua H, Braun M-G, Burdick DJ, Castanedo GM, Chen Y-C, Cheng Y-X, Cheong J, Daniels B, Deshmukh G, Fu Y, Gibbons P, Gloor SL, Hua R, Labadie S, Liu X, Pastor R, Stivala C, Xu M, Xu Y, Zheng H, Kapadia SB, Hanan EJ. 2020. Optimization of globomycin analogs as novel gram-negative antibiotics. Bioorg Med Chem Lett 30:127419.

39. Diao J, Komura R, Sano T, Pantua H, Storek KM, Inaba H, Ogawa H, Noland CL, Peng Y, Gloor SL, Yan D, Kang J, Katakam AK, Volny M, Liu P, Nickerson NN, Sandoval W, Austin CD, Murray J, Rutherford ST, Reichelt M, Xu Y, Xu M, Yanagida H, Nishikawa J, Reid PC, Cunningham CN, Kapadia SB. 2021. Inhibition of Escherichia coli Lipoprotein Diacylglyceryl Transferase Is Insensitive to Resistance Caused by Deletion of Braun’s Lipoprotein. J Bacteriol 203:e00149–21.

40. Huang K-J, Pantua H, Diao J, Skippington E, Volny M, Sandoval W, Tiku V, Peng Y, Sagolla M, Yan D, Kang J, Katakam AK, Michaelian N, Reichelt M, Tan M-W, Austin CD, Xu M, Hanan E, Kapadia SB. 2022. Deletion of a previously uncharacterized lipoprotein lirL confers resistance to an inhibitor of type II signal peptidase in Acinetobacter baumannii. Proc National Acad Sci 119:e2123117119.

41. Phan M-D, Peters KM, Sarkar S, Lukowski SW, Allsopp LP, Moriel DG, Achard MES, Totsika M, Marshall VM, Upton M, Beatson SA, Schembri MA. 2013. The Serum Resistome of a Globally Disseminated Multidrug Resistant Uropathogenic Escherichia coli Clone. PLoS Genet 9:e1003834.

42. Sha J, Fadl AA, Klimpel GR, Niesel DW, Popov VL, Chopra AK. 2004. The Two Murein Lipoproteins of Salmonella enterica Serovar Typhimurium Contribute to the Virulence of the Organism. Infect Immun 72:3987–4003.

43. Fadl AA, Sha J, Klimpel GR, Olano JP, Niesel DW, Chopra AK. 2005. Murein Lipoprotein Is a Critical Outer Membrane Component Involved in Salmonella enterica Serovar Typhimurium Systemic Infection. Infect Immun 73:1081–1096.

44. Sha J, Agar SL, Baze WB, Olano JP, Fadl AA, Erova TE, Wang S, Foltz SM, Suarez G, Motin VL, Chauhan S, Klimpel GR, Peterson JW, Chopra AK. 2008. Braun Lipoprotein (Lpp) Contributes to Virulence of Yersiniae: Potential Role of Lpp in Inducing Bubonic and Pneumonic Plague. Infect Immun 76:1390–1409.

45. Liu T, Agar SL, Sha J, Chopra AK. 2010. Deletion of Braun lipoprotein gene (lpp) attenuates Yersinia pestis KIM/D27 strain: Role of Lpp in modulating host immune response, NF-κB activation and cell death. Microb Pathog 48:42–52.

46. Pantua H, Skippington E, Braun M-G, Noland CL, Diao J, Peng Y, Gloor SL, Yan D, Kang J, Katakam AK, Reeder J, Castanedo GM, Garland K, Komuves L, Sagolla M, Austin CD, Murray J, Xu Y, Modrusan Z, Xu M, Hanan EJ, Kapadia SB. 2020. Unstable Mechanisms of Resistance to Inhibitors of Escherichia coli Lipoprotein Signal Peptidase. mBio 11:e02018–20.

47. Bei W, Luo Q, Shi H, Zhou H, Zhou M, Zhang X, Huang Y. 2022. Cryo-EM structures of LolCDE reveal the molecular mechanism of bacterial lipoprotein sorting in Escherichia coli. Plos Biol 20:e3001823.

48. Sharma S, Zhou R, Wan L, Feng S, Song K, Xu C, Li Y, Liao M. 2021. Mechanism of LolCDE as a molecular extruder of bacterial triacylated lipoproteins. Nat Commun 12:4687.

49. Okuda S, Watanabe S, Tokuda H. 2008. A short helix in the C-terminal region of LolA is important for the specific membrane localization of lipoproteins. FEBS letters 582:2247–2251.

50. Narita S, Tokuda H. 2011. Overexpression of LolCDE Allows Deletion of the Escherichia coli Gene Encoding Apolipoprotein N-Acyltransferase. J Bacteriol 193:4832– 4840.

51. Smith HC, May KL, Grabowicz M. 2023. Teasing apart the evolution of lipoprotein trafficking in gram-negative bacteria reveals a bifunctional LolA. Proc National Acad Sci 120:e2218473120.

52. Hayashi Y, Tsurumizu R, Tsukahara J, Takeda K, Narita S, Mori M, Miki K, Tokuda H. 2014. Roles of the Protruding Loop of Factor B Essential for the Localization of Lipoproteins (LolB) in the Anchoring of Bacterial Triacylated Proteins to the Outer Membrane. J Biol Chem 289:10530–10539.

53. Baba T, Ara T, Hasegawa M, Takai Y, Okumura Y, Baba M, Datsenko KA, Tomita M, Wanner BL, Mori H. 2006. Construction of Escherichia coli K-12 in-frame, single-gene knockout mutants: the Keio collection. Mol Syst Biology 2:2006.0008-2006.0008.

54. Datsenko KA, Wanner BL. 2000. One-step inactivation of chromosomal genes in Escherichia coli K-12 using PCR products. Proc National Acad Sci 97:6640–6645.

55. Nichols BP, Shafiq O, Meiners V. 1998. Sequence Analysis of Tn 10 Insertion Sites in a Collection of Escherichia coli Strains Used for Genetic Mapping and Strain Construction. J Bacteriol 180:6408–6411.

56. Yu D, Ellis HM, Lee E-C, Jenkins NA, Copeland NG, Court DL. 2000. An efficient recombination system for chromosome engineering in Escherichia coli. Proc National Acad Sci 97:5978–5983.

57. Shrivastava R, Jiang X, Chng S. 2017. Outer membrane lipid homeostasis via retrograde phospholipid transport in Escherichia coli. Mol Microbiol 106:395–408.

58. Douglass MV, Cléon F, Trent MS. 2021. Cardiolipin aids in lipopolysaccharide transport to the gram-negative outer membrane. Proc Natl Acad Sci 118:e2018329118.

59. Wu T, McCandlish AC, Gronenberg LS, Chng S-S, Silhavy TJ, Kahne D. 2006. Identification of a protein complex that assembles lipopolysaccharide in the outer membrane of Escherichia coli. Proc Natl Acad Sci 103:11754–11759.

60. Teufel F, Armenteros JJA, Johansen AR, Gíslason MH, Pihl SI, Tsirigos KD, Winther O, Brunak S, Heijne G von, Nielsen H. 2022. SignalP 6.0 predicts all five types of signal peptides using protein language models. Nat Biotechnol 40:1023–1025.

