## Supplemental Material for "Genetic analysis reveals a robust and hierarchical recruitment of the LolA chaperone to the LolCDE lipoprotein transporter"

```
PDB   -------------------------------------DAASDLKSRLDKVSSFHASFTQK   23
Eco   -------------------------------------DAASDLKSRLDKVSSFHASFTQK   23
Vch   -------------------------------------APKDTLSERLAMSEGFSATFNQQ   23
Pae   ----------------------------------DDSAAVQRLTGLLNKAQTLTARFSQL   26
Lpn   ------------------------------------QTSAEVLQSKLNAIQTMTANFSQI   24
Bpe   ------------------------------------ATAQEQLDTFVATVKGATGSFKQS   24
Bps   -------------------------------------SGTEQLKAFVAKVHAAKGEFTQQ   23
Ccr   -------------------QGSTAPVPAQLSAEQKA--LLDKATAYIQGLSSAKGRFVQT   39
Bms   KAGRVAAVLGMGILGVGVAGAALAPVPVKAQGADGGIAAAQQIADHFSSARTMTGEFVQF   60
Nme   -----------------------------------QAGAVDALKQFNNDADGISGSFTQT   25
Cje   ---------------------------------------------LDLNFNTFSSNFTQI   15
Hpy   ----------------------------------KNPSTLSKEEEVLQHLQSFSAHFKQV   26
                                                            . * *

PDB   VTDG----------SGAAVQEGQGDLWVKRPNLFNWHMTQPDESILVSDGKTLWFYNPFV   73
Eco   VTDG----------SGAAVQEGQGDLWVKRPNLFNWHMTQPDESILVSDGKTLWFYNPFV   73
Vch   VLSP----------EGKVILTGNGKVDIARPSLFRWETETPDENLLVSDGTTLWHFDPFV   73
Pae   TLDG----------SGTRLQETAGQLSLKRPGLFRWHTDAPNEQLLISNGEKVWLYDPDL   76
Lpn   VK-A----------KNREVSRSSGSMALQRPGRFRWQTKDPLEQLIVADGQKMWIYDVDL   73
Bpe   TVSP----------QGATQPAQSGTFAFQRPGKFKWAVQLPYEQLIVSDGKQVFQYDPDL   74
Bps   IVKAPPKAASGAVPTVKPTDNSSGSFVFSRPGKFIWTYQKPYQQVLQADGDKLYVYDKDL   83
Nme   ------------VQSKKKTQTAHGTFKILRPGLFKWEYTKPYRQTIVGDGQTVWLYDVDL   73
Ccr   DQR---------------GTQTQGTFYLQRPGKARFAYDPPAGLLVVSNGNNVNIFDSRL   84
Bms   GPR---------------GEQTGGTFYIERPGKIRFNYNNSP-IRVISDGDSVVINNRKL   104
Cje   VKSK------------N-STLSYSGHFILSKDQAYWSYDTPSKKEIYINKNQVTIVEHDL   62
Hpy   LKNE------------K-PLVYYGVLKAKAPNWALWVYEKPLKKEIYMNDKEVVIYEPNL   73
                             .       .   :         :  :   :   :  :

PDB   EQATATWLKDATGNTPFMLIARNQSSDWQQYNIKQNG-----DDFVLTP-KASNGNLKQF   127
Eco   EQATATWLKDATGNTPFMLIARNQSSDWQQYNIKQNG-----DDFVLTP-KASNGNLKQF   127
Vch   EQVTLYRAEEALEQTPFVLLTRNKASDWDAYHVEEKG-----DVFTLTP-TALDSNQGRF   127
Pae   EQVTIQKLDQRLTQTPALLLSGDISKISESFAITYK-EGGNVVDFVLKP-KTKDTLFDTL   134
Lpn   EQVTVKNQEKGLGGTAALFLSGYDETLTHDFDVSEK-QKGKLTVFDLKS-KSAKENFQRI   131
Bpe   AQVTVRQVDQAIGTSPAAILFGAGQ-LSQAFAVSALPDRDGLQWLRAKP-RNADAGFSQV   132
Bps   NQVTERKLAGALGASPAAILFGSND-LDKNYTLRDAGEKGGIDWLEMVP-KAQDTQFQRI   141
Nme   AQVTKSSQDQAIGGSPAAILSNKTA-LESSYTLKEDGSSNGIDYVLATP-KRNNAGYQYI   131
Ccr   KTYESYPLSK----TPLNLLLAREVRLDRGVVITDVR--PLADGFTIVAQDAKRQALGRI   138
Bms   DTWDLYPLSK----TPLKLLLADRIDLGGGRLQS-VK--QEPDMTTLVLGDKSVFGDSKI   157
Cje   EQVIFSHLDNIPNLN--EIFKKASLI----------------DKDKLVA-KYDN---INY   100
Hpy   FQATITPLKDKTDFF--TILKRLKKQ----------------DDGSFKT-TINK---TTY   111
                        ::

PDB   TINVGRDGT--IHQFSAVEQDDQRSSYQLKSQQNG-AVDAAKFTFTP-PQGVTVDDQRK   182
Eco   TINVGRDGT--IHQFSAVEQDDQRSSYQLKSQQNG-AVDAAKFTFTP-PQGVTVDDQRK   182
Vch   QITISEKGV--VQGFKVIEQDGQQSEFTFSKVKQQ-KPNASVFNYKV-PKGVEVDDQRN   182
Pae   RLSF-RSGK--VNDMQMIDGVGQRTNILFFDVKMNEALDAKQFTFDV-PPGVDVIQE--   187
Lpn   KLIF-SQST--LIGLELYDQLGQITDVKLVQIKSNPKLPAKLFQFKP-PKGVDVVKQ--   184
Bpe   DIGL-RDNQ--PARIELVDAFGQTTRVELSNLLPG-AVPASEFQFTP-PQGVDVVKM--   184
Bps   GIGF-RNGM--LAAMELHDVFGNVTLLTFTNIQTNPPLKADQFKFVV-PKGADVVTG--   194
Nme   RIGF-KGGN--LAAMQLKDSFGNQTSISFGGLNTNPQLSRGAFKFTP-PKGVDVLSN--   184
Ccr   SIDF-SNGLVGLMGWTVTDIKGGQIRVRLSDFAETADLDPKLFVLTD-PRRKVGKP---   192
Bms   TMMF-DPKSYDLKQWTITDAQKLDTTVMIFNVRTGVRFTNDMFKIDY-QRIAMKRKGQ-   213
Cje   TIKL-NQEQ--IQSISYKDEFENDVIINLNNQIKNPKINSDVFKAKI-PQNYDIVR---   152
Hpy   RLVF-KDGK--PFSLEFKDGMNNLVTITFSQAEINPTIANEIFVFKPKDENIDIVRQ--   165
       : .              :         :             *
```

**Fig. S1. Predicted LolA sites that make polar contacts with the LolC Pad.** Aligned mature LolA protein sequences with yellow highlighted residues that are predicted by AlphaFold2 Multimer to make polar contacts to their native LolC or LolF. Blue highlighting indicates low confidence prediction. PDB denotes structure 6F3Z. LolA sequences are from *Escherichia coli* MG1655 (*Eco*), *Vibrio cholerae* O1 El Tor N16961 (*Vch*),

*Pseudomonas aeruginosa* PAO1 (*Pae*), *Legionella pneumophila* subsp. pneumophila Philadelphia 1 (*Lpn*), *Bordetella pertussis* Tohama I (*Bpe*), *Burkholderia pseudomallei* K96243 (*Bps*), *Neisseria meningitidis* MC58 (*Nme*), *Caulobacter vibrioides* CB15 (*Ccr*), *Brucella suis* 1330 (Bms), *Campylobacter jejuni* subsp. jejuni NCTC 11168 (*Cje*), *Helicobacter pylori* 26695 (*Hpy*). Signal peptide processed mature LolA sequences were predicted with SignalP-6.0. Alignments were made with Clustal Omega.

```
                                                   |E.coli Hook|
PDB VKQTDLEPGKYN---VILGEQLASQLGVNRGDQIRVMVPSASQFTPMGRIPSQRLFNVIG 188
Eco VKQTDLEPGKYN---VILGEQLASQLGVNRGDQIRVMVPSASQFTPMGRIPSQRLFNVIG 188
Vch GRVTALQAGEYQ---LFLGHLLARSLNVTVGDKVRLMVTEASQFTPLGRLPSQRNFTVAG 193
Pae GSLDDLKPGEFG---IVLGEITARRFHVNVGDKLTLIVPEAT-SAPGGITPRMQRFTIVA 209
Lpn GNMSNLK--HFG---IILGKGLADSLGVMIGDKVTIMIPQAT-VTPAGMIPRFKRFTVVG 225
Bpe GKLSDLVAGGFG---AVLGSDLADGLGVKTGDTVLMLAPQGS-ISPAGFAPRMRQFTVVG 187
Bps GALTALAPGQFG---IVLGNALAGNLGVGVGDKVTLVAPEGT-ITPAGMMPRLKQFTVVG 194
Nme GKFEDLIPGEFD---IILGVGLAEALGAEVGNKVTVITPEGN-VTPAGVVPRLKQFTVVG 190
Ccr GSMEGFGQGEYGGDIVLIGERMAQTLGVQPGDPITIISPSGP-ATAFGSSTREKNYIVGG 202
Bms GTLKGFDHS----GGVAIGTRMAENLGLSVGDTLRVISPDGD-VTPFGVNPRVKAYPIVA 198
Cje ENL-----SGFD---ILVGSALTDEFGLHKNDKLSLIFSNLN-PSGFSLVPQTKRFDVKA 186
Hpy INENDLFKNPFN---LIVGKSLRYSLNLDLNQKADLFFTELE-PTGLTLSPIMKRFTIKG 192
                     :*      :    .:   :.  .    :        : : : .
```

**Fig. S2. LolC/LolF Pad sites predicted to make polar contacts with LolA.** Aligned mature LolC or LolF protein sequences from the region spanning the Pad. Yellow highlighted residues are predicted by AlphaFold2 Multimer to make polar contacts to their native LolA. Blue highlighting indicates low confidence prediction. PDB denotes structure 6F3Z. LolC sequences are from *Escherichia coli* MG1655 (*Eco*), *Vibrio cholerae* O1 El Tor N16961 (*Vch*), *Pseudomonas aeruginosa* PAO1 (*Pae*). LolF sequences are from *Legionella pneumophila* subsp. pneumophila Philadelphia 1 (*Lpn*), *Bordetella pertussis* Tohama I (*Bpe*), *Burkholderia pseudomallei* K96243 (*Bps*), *Neisseria meningitidis* MC58 (*Nme*), *Caulobacter vibrioides* CB15 (*Ccr*), *Brucella suis* 1330 (Bms), *Campylobacter jejuni* subsp. jejuni NCTC 11168 (*Cje*), *Helicobacter pylori* 26695 (*Hpy*). Alignments were made with Clustal Omega.

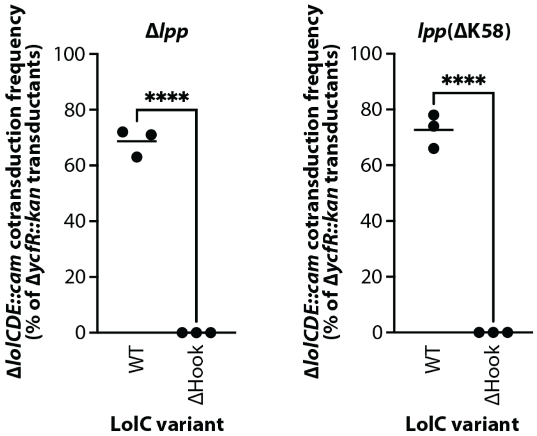


**Figure S3. The LolC Hook is essential even when Lpp is detoxified or entirely absent.** Genetic linkage between Δ*ycfR*::*kan* and Δ*lolCDE*::*cam* when introduced into *E. coli* mutants either lacking *lpp* or the Lpp K58 residue and carrying pBAD18::*lolCDE* plasmids encoding *lolC*(ΔHook) mutations. Three independent transductions were performed and 100 transductants from each experiment where tested (n=100 per data point); statistically significant results were found by one-way Anova (**** indicates $P$<0.0001).

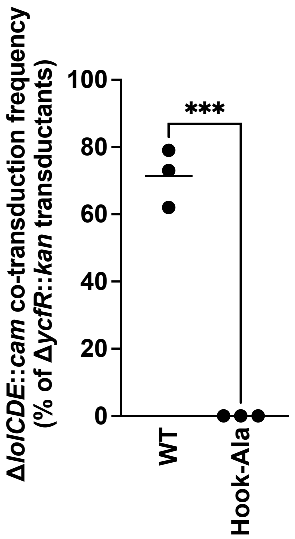


**Figure S4. Plasmid expressed *lolC*(Hook-Ala) is unable to complement loss of chromosomal *lolC* even in Δ*lpp E. coli.*** Genetic linkage between Δ*ycfR*::*kan* and Δ*lolCDE*::*cam* when introduced into *E. coli* mutants lacking *lpp* and carrying pBAD18::*lolCDE* plasmids encoding *lolC*(Hook-Ala) or *lolC*+. . Three independent transductions were performed and 100 transductants from each experiment where tested (n=100 per data point); statistically significant results were found by unpaired t-test (**** indicates *P*<0.0001).

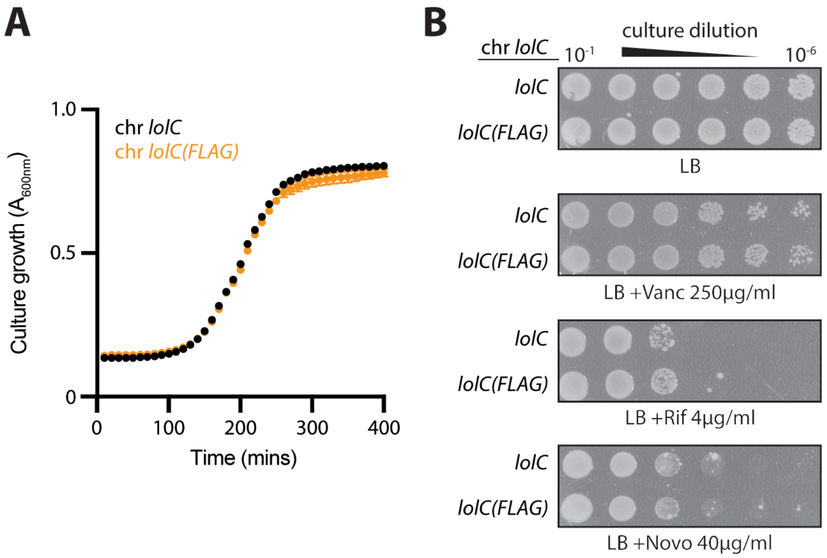


**Figure S5. Appending an N-terminal FLAG tag to LolC does not appreciably affect LolC function.** DNA sequence encoding an N-terminal FLAG tag was introduced into the native *lolC* locus. **(A)** Growth of a chromosomal *lolC(FLAG)* or wildtype *lolC*+ was assessed at 37°C by measuring culture turbidity ($A_{600nm}$) over time. **(B)** OM antibiotic barrier function was assessed by testing antibiotic sensitivity at 37°C with the antibiotics indicated.

**Table S1: Strains used in this study**

**Table S2 Plasmids used in this study**

**Table S3: Oligonucleotides used in this study**

**Table S1. Strains used in this study**

| Name | Genotype | Reference |
|---|---|---|
| MC4100 | MC4100 F- {[araD139]B/r Δ(argF-lac)169 λ- e14- flhD5301 Δ(fruK-yeiR)725(fruA25) relA1 rpsL150 rbsR22 thi-1 Δ(fimB-fimE)632(::IS1) deoC1} | (1) |
| NR756 | MC4100 Ara$^+$ | (2) |
| DY378 | W3110 λcI857 Δ(*cro-bioA*) | (3) |
| KL351 | MC4100 Ara$^+$ [pBAD18::*lolCDE*] | (4) |
| KL379 | MC4100 Ara$^+$ [pBAD18::*lolC*(ΔHook)*DE*] | This study |
| KM999 | MC4100 Ara$^+$ [pBAD18::*lolC*(M175R)*DE*] | This study |
| KL524 | MC4100 Ara$^+$ [pBAD18::*lolC*(S168A)*DE*] | This study |
| KL515 | MC4100 Ara$^+$ [pBAD18::*lolC*(S168D)*DE*] | This study |
| KL549 | MC4100 Ara$^+$ [pBAD18::*lolC*(S168R)*DE*] | This study |
| KL540 | MC4100 Ara$^+$ [pBAD18::*lolC*(S170A)*DE*] | This study |
| KL552 | MC4100 Ara$^+$ [pBAD18::*lolC*(S170D)*DE*] | This study |
| KL527 | MC4100 Ara$^+$ [pBAD18::*lolC*(S170R)*DE*] | This study |
| KL521 | MC4100 Ara$^+$ [pBAD18::*lolC*(Q171A)*DE*] | This study |
| KL390 | MC4100 Ara$^+$ [pBAD18::*lolC*(T173A)*DE*] | This study |
| KL518 | MC4100 Ara$^+$ [pBAD18::*lolC*(R177A)*DE*] | This study |
| KL530 | MC4100 Ara$^+$ [pBAD18::*lolC*(R177D)*DE*] | This study |
| KL460 | MC4100 Ara$^+$ [pBAD18::*lolC*(I178A)*DE*] | This study |
| MG3437 | MC4100 Ara$^+$ Δ*ycfR*::*kan* Δ*lolCDE*::*cam* | This study |
| KL466 | MC4100 Ara$^+$ Δ*lolCDE*::*cam* [pBAD18::*lolCDE*] | This study |
| KM1001 | MC4100 Ara$^+$ Δ*lolCDE*::*cam* [pBAD18::*lolC*(M175R)*DE*] | This study |
| KL538 | MC4100 Ara$^+$ Δ*lolCDE*::*cam* [pBAD18::*lolC*(S168A)*DE*] | This study |
| KL534 | MC4100 Ara$^+$ Δ*lolCDE*::*cam* [pBAD18::*lolC*(S168D)*DE*] | This study |
| KL567 | MC4100 Ara$^+$ Δ*lolCDE*::*cam* [pBAD18::*lolC*(S168R)*DE*] | This study |
| KL565 | MC4100 Ara$^+$ Δ*lolCDE*::*cam* [pBAD18::*lolC*(S170A)*DE*] | This study |
| KL569 | MC4100 Ara$^+$ Δ*lolCDE*::*cam* [pBAD18::*lolC*(S170D)*DE*] | This study |
| KL558 | MC4100 Ara$^+$ Δ*lolCDE*::*cam* [pBAD18::*lolC*(S170R)*DE*] | This study |
| KL537 | MC4100 Ara$^+$ Δ*lolCDE*::cam [pBAD18::*lolC*(Q171A)*DE*] | This study |
| KL463 | MC4100 Ara$^+$ Δ*lolCDE*::cam [pBAD18::*lolC*(T173A)*DE*] | This study |
| KL535 | MC4100 Ara$^+$ Δ*lolCDE*::cam [pBAD18::*lolC*(R177A)*DE*] | This study |
| KL561 | MC4100 Ara$^+$ Δ*lolCDE*::cam [pBAD18::*lolC*(R177D)*DE*] | This study |
| KL493 | MC4100 Ara$^+$ Δ*lolCDE*::cam [pBAD18::*lolC*(I178A)*DE*] | This study |
| KL980 | MC4100 Ara$^+$ [pBAD18::FLAG-*lolCDE*] | This study |
| KL1178 | MC4100 Ara$^+$ [pBAD18::FLAG-*lolC*(ΔHook)*DE*] | This study |
| KL1237 | MC4100 Ara$^+$ [pBAD18::FLAG-*lolC*(ΔM175R)*DE*] | This study |
| MG4583 | MC4100 Ara$^+$ [pBAD18::FLAG-*lolCDE*] | This study |
| KL1272 | MC4100 Ara$^+$ [pBAD18::*lolC*(R163A)*DE*] | This study |
| KM987 | MC4100 Ara$^+$ [pBAD18::*lolC*(Q181A)*DE*] | This study |
| KM988 | MC4100 Ara$^+$ [pBAD18::*lolC*(R163A Q181A)*DE*] | This study |
| KM989 | MC4100 Ara$^+$ [pBAD18::*lolC*(R182A)*DE*] | This study |
| KM990 | MC4100 Ara$^+$ [pBAD18::*lolC*(R163A R182A)*DE*] | This study |
| KM991 | MC4100 Ara$^+$ [pBAD18::*lolC*(Q181A R182A)*DE*] | This study |
| KM992 | MC4100 Ara$^+$ [pBAD18::*lolC*(R163A Q181A R182A)*DE*] | This study |

| | | |
|---|---|---|
| KM799 | MC4100 Ara+ Δ*lolCDE*::cam [pBAD18::*lolCDE*] | This study |
| KM993 | MC4100 Ara+ Δ*lolCDE*::cam [pBAD18::*lolC*(R163A)*DE*] | This study |
| KL537 | MC4100 Ara+ Δ*lolCDE*::cam [pBAD18::*lolC*(Q181A)*DE*] | This study |
| KM994 | MC4100 Ara+ Δ*lolCDE*::cam [pBAD18::*lolC*(R163A Q181A)*DE*] | This study |
| KL464 | MC4100 Ara+ Δ*lolCDE*::cam [pBAD18::*lolC*(R182A)*DE*] | This study |
| KM995 | MC4100 Ara+ Δ*lolCDE*::cam [pBAD18::*lolC*(R163A R182A)*DE*] | This study |
| KM996 | MC4100 Ara+ [pBAD18::*lolC*(Q181A R182A)*DE*] | This study |
| KM997 | MC4100 Ara+ [pBAD18::*lolC*(R163A Q181A R182A)*DE*] | This study |
| KM1002 | MC4100 Ara+ *zce-726*::Tn*10 lolC+* | This study |
| MG4644 | MC4100 Ara+ *zce-726*::Tn*10 FLAG-lolC+* | This study |
| KM1003 | MC4100 Ara+ *zce-726*::Tn*10* FLAG-*lolC*(R163A) | This study |
| KM1004 | MC4100 Ara+ *zce-726*::Tn*10* FLAG-*lolC*(Q181A R182A) | This study |
| KM998 | MC4100 Ara+ *zce-726*::Tn*10* FLAG-*lolC*(R163A Q181A R182A) [pBAD18::*lolCDE*] | This study |
| KM1005 | MC4100 Ara+ Δ*ynhG lpp*(ΔK58) *zce-726*::Tn*10* FLAG-*lolC* | This study |
| KM1006 | MC4100 Ara+ Δ*ynhG lpp*(ΔK58) *zce-726*::Tn*10* FLAG- *lolC*(R163A) | This study |
| KM1007 | MC4100 Ara+ Δ*ynhG lpp*(ΔK58) *zce-726*::Tn*10* FLAG- *lolC*(Q181A R182A) | This study |
| KM1008 | MC4100 Ara+ Δ*ynhG lpp*(ΔK58) *zce-726*::Tn*10* FLAG-*lolC* (R163A Q181A R182A) | This study |
| KM1014 | MC4100 Ara+ *zce-726*::Tn*10* FLAG-*lolC*(M175R) | This study |
| KM1015 | MC4100 Ara+ Δ*ynhG lpp*(ΔK58) *zce-726*::Tn*10* FLAG-*lolC*(M175R) | This study |
| KM1017 | MC4100 Ara+ Δ*lpp* [pBAD18::*lolC*(ΔHook)*DE*] | This study |
| MG4726 | MC4100 Ara+ [pBAD18::*lolC*(T173A M175R I178A)*DE*] | This study |
| MG4702 | MC4100 Ara+ [pBAD18::FLAG-*lolC*(T173A M175R I178A)*DE*] | This study |
| MG4727 | MC4100 Ara+ [pBAD18::*lolC*(Hook-Ala)*DE*] | This study |
| MG4705 | MC4100 Ara+ [pBAD18::FLAG-*lolC*(Hook-Ala)*DE*] | This study |
| MG4728 | MC4100 Ara+ Δ*lpp* [pBAD18::-*lolC*(Hook-Ala)*DE*] | This study |
| KM1018 | MC4100 Ara+ *lpp*(ΔK58) [pBAD18::*lolC*(ΔHook)*DE*] | This study |
| MG4595 | DY378 Δ*lpp*::*kan* Δ*lolCDE*::*cam* [pBAD18::*lolCDE*] | This study |
| MG2502 | MC4100 Ara+ Δ*ynhG lpp*(ΔK58) *zce-726*::Tn*10* | This study |
| MG4655 | MC4100 Ara+ Δ*ynhG lpp*(ΔK58) *zce-726*::Tn*10 lolC* (R163A Q181A R182A) | This study |

1. Casadaban MJ. 1976. Transposition and fusion of the lac genes to selected promoters in Escherichia coli using bacteriophage lambda and Mu. J Mol Biology 104:541–555.
2. Button JE, Silhavy TJ, Ruiz N. 2007. A Suppressor of Cell Death Caused by the Loss of σ E Downregulates Extracytoplasmic Stress Responses and Outer Membrane Vesicle Production in Escherichia coli. J Bacteriol 189:1523–1530.
3. Yu D, Ellis HM, Lee E-C, Jenkins NA, Copeland NG, Court DL. 2000. An efficient recombination system for chromosome engineering in Escherichia coli. Proc National Acad Sci 97:5978–5983.
4. Grabowicz M, Silhavy TJ. 2017. Redefining the essential trafficking pathway for outer membrane lipoproteins. Proc National Acad Sci 114:4769–4774.

**Table S2. Plasmids used in this study**

| Name | Description | Reference |
|---|---|---|
| pBAD18 | Arabinose inducible, $Amp^R$, cloning vector | (1) |
| pABD18::lolCDE | *lolCDE* cloned to be arabinose inducible, $Amp^R$ | (2) |

1. Guzman LM, Belin D, Carson MJ, Beckwith J. 1995. Tight regulation, modulation, and high-level expression by vectors containing the arabinose PBAD promoter. J Bacteriol 177:4121–4130.
2. Grabowicz M, Silhavy TJ. 2017. Redefining the essential trafficking pathway for outer membrane lipoproteins. Proc National Acad Sci 114:4769–4774.

**Table S3. Oligonucleotides used in this study**

| Name | Sequence | Use |
|---|---|---|
| deltahookLolC _F | ccaagccagcgcctgttc | *lolC* mutagenesis |
| deltahookLolC _R | tggtaccatcacgcggatttg | *lolC* mutagenesis |
| hook-Ala-LolC _F | gcgccggcggcggctgctccaagccagcgc | *lolC* mutagenesis |
| hook-Ala-LolC _R | ggccgcggcggcagctggtaccatcacgcg | *lolC* mutagenesis |
| T173A M175R I178A LolC _F | gcgccgcgtgggcgtgctccaagccagcgc | *lolC* mutagenesis |
| T173A M175R I178A LolC _R | gaactggctggcagat | *lolC* mutagenesis |
| R163A_LolC_ R | ccgcgattaacgcctagc | *lolC* mutagenesis |
| R163A_LolC_ F | tgatcaaatcgccgtgatggtaccatctgc | *lolC* mutagenesis |
| M175R_LolC_ F | gttcacgccgcgtgggcgtattccaagc | *lolC* mutagenesis |
| M175R_LolC_ R | tggctggcagatggtacc | *lolC* mutagenesis |
| T173A_LolC_ F | cagccagttcgctccgatggggc | *lolC* mutagenesis |
| T173A_LolC_ R | gcagatggtaccatcacgc | *lolC* mutagenesis |
| I178A_LolC_F | gatggggcgtgctccaagccagcg | *lolC* mutagenesis |
| R182A_LolC_ F | tccaagccaggccctgttcaatgtgattggc | *lolC* mutagenesis |
| 182A_LolC_R | atacgccccatcggcgtg | *lolC* mutagenesis |
| I178A_LolC_R | ggcgtgaactggctggca | *lolC* mutagenesis |
| Q181A_LolC_ R | tattccaagcgcgcgcctgttcaatgtgattggc | *lolC* mutagenesis |
| Q181A_LolC_ F | cgccccatcggcgtgaac | *lolC* mutagenesis |
| Q171A_LolC_ F | atctgccagcgcgttcacgccgatg | *lolC* mutagenesis |
| Q171A_LolC_ R | ggtaccatcacgcggatt | *lolC* mutagenesis |
| R177A_LolC_ R | gtgaactggctggcagat | *lolC* mutagenesis |
| R177A_LolC_ F | gccgatgggggctattccaagccag | *lolC* mutagenesis |
| F172A_LolC_ R | gatggtaccatcacgcgg | *lolC* mutagenesis |
| F172A_LolC_F | tgccagccaggccacgccgatgg | *lolC* mutagenesis |
| S168A_F | gcggccagccagttcac | *lolC* mutagenesis |
| S168_R | tggtaccatcacgcggatttg | *lolC* mutagenesis |
| S168D_F | gatgccagccagttcacg | *lolC* mutagenesis |
| S168R_F | cgtgccagccagttcac | *lolC* mutagenesis |
| S170A_F | gcgcagttcacgccgatg | *lolC* mutagenesis |

| | | |
|---|---|---|
| S170_R | ggcagatggtaccatcac | *lolC* mutagenesis |
| S170D_F | gaccagttcacgccgatg | *lolC* mutagenesis |
| S170R_F | cgtcagttcacgccgatg | *lolC* mutagenesis |
| Q171A_F | gcgttcacgccgatgggg | *lolC* mutagenesis |
| Q171_R | gctggcagatggtaccatcac | *lolC* mutagenesis |
| R177A_F | gctattccaagccagcgc | *lolC* mutagenesis |
| R177_R | ccccatcggcgtgaactg | *lolC* mutagenesis |
| R177D_F | gatattccaagccagcgc | *lolC* mutagenesis |
| F172_R | ctggctggcagatggtac | *lolC* mutagenesis |
| F172S_F | tctacgccgatggggc | *lolC* mutagenesis |
| F172R_R | cgtacgccgatgggg | *lolC* mutagenesis |
| F172D_F | gatacgccgatggggcg | *lolC* mutagenesis |
| LolC_1812_R | gcttggaatacgcccc | *lolC* mutagenesis |
| LolC_Q181A_F | gcgcgcctgttcaatgtgatt | *lolC* mutagenesis |
| LolC_R182A_F | caggcgctgttcaatgtgattggcactttc | *lolC* mutagenesis |
| LolC_QR181AA_F | gcggcgctgttcaatgtgattggcactttc | *lolC* mutagenesis |
| rec_p18lolCDE_fwd | gattgatttacgggggcttttcagattagccctgacgatcacttacagttcagacgtttacccat cttgctttcgcttatatactcgtgtctttgctacagcaacc | Recombineering template |
| rec_p18lolCDE_rev | ctgcaactgccgaccgctatcaaacacgccaagcgcaatttttgttccaccaatatcaaacccgt aatacattgccgctccttgttttaatgtactgcctttactgg | Recombineering template |
| KM178LolC_N -FLAG_F | atggactacaaagacgatgacgacaagtaccaacctgt cgctctatt | FLAG tagging |
| KM177LolC_N -tag_R | gaaatccgtctggttgctgtag | FLAG tagging |
